# The exit of naïve pluripotency contains a lipid metabolism-induced checkpoint for genome integrity

**DOI:** 10.1101/2024.09.22.613751

**Authors:** Roshni A. de Souza, David Barneda, Donja Karimlou, Nick G.P. Bovee, Yuhan Zheng, Eveline J.E.M. Kahlman, Clara Lopes Novo, James K. Ellis, Bryony J. Leeke, Megha Prakash Bangalore, Zijing Liu, Bebiana C. Sousa, Andrea F. Lopez-Clavijo, Joop H. Jansen, Mauricio Barahona, Michelle Percharde, Hector C. Keun, Mark Christian, Hendrik Marks, Véronique Azuara

## Abstract

Pluripotent progenitors undergo dramatic cellular and biochemical transformations during peri-implantation development. These large-scale reprogramming events are fundamental for subsequent differentiation, but how they are integrated and co-ordinated with the preservation of genome integrity remain unknown. Here, we uncover a metabolism-induced telomere checkpoint that takes place in pluripotent progenitors as they form rosette-like epithelial structures. We show that the glycolytic switch at the exit of naïve pluripotency is preceded by an acceleration of mitochondrial respiration and *de novo* lipogenesis, fuelling the accumulation of lipid droplets required for morphogenesis. We find that downstream of these CIDEA-promoted metabolic events is the induction of ZSCAN4, a key pluripotency-associated regulator of telomere stability. Surprisingly, the build-up of lipid droplets corresponds to a transient shortening of telomeres, which triggers the activation of an elongation mechanism via ZSCAN4. Thus, telomere homeostasis can be safeguarded as essential lipid metabolic reprogramming unfolds to drive developmental progression.

## Introduction

Mammalian life begins with the fusion of an oocyte and a sperm to form a zygote. Through ensuing cleavage divisions, the embryo undergoes a series of biochemical changes and cell state transitions to form the blastocyst^1^. Preserving the genome integrity of cells while going through extensive epigenomic and metabolic reprogramming is critical for the early embryo, given its potential to generate all somatic cell types of an adult organism^2^. The development of pluripotent progenitors, which first emerge within the inner cell mass of the blastocyst, provides a fundamentally important paradigm for understanding how cell state transitions are executed while safeguarding the developmental capacity of cells. At peri-implantation, pluripotent progenitors mature from a naïve to a more developmentally primed state in preparation for differentiation^3^. This transition occurs as cells form rosette-like epithelial structures around an expanding lumen, a prerequisite for all subsequent development *in vivo*^4,5^. Naïve and primed progenitors can also be captured and propagated *in vitro* as embryonic stem cells (ESCs) and epiblast stem cells (EpiSCs), respectively^6–9^. These stem cell populations have been instrumental to delineate the cellular, biochemical and molecular features that distinguish pluripotent cell states^8–14^. More recently, three-dimensional (3D) spheroid models of ESC differentiation have enabled further investigations of pluripotency maturation in a tissue context^5,15^. Yet, we still have little understanding of how the different reprogramming events that underline this transition converge and are coordinated to support developmental progression upon implantation.

Metabolic remodelling is hardwired into the complex programme of early embryo development^16–20^. Previous studies indicated that the newly formed blastocyst accumulates a substantial amount of stored lipids (e.g., triglycerides) in the form of lipid droplets (LDs), which are then mobilized upon implantation^17,21–25^. Remarkably, this dynamic behaviour of LDs plays a key role in orchestrating the cellular remodelling of pluripotent stem cells, highlighting a surprising interconnection between lipid storage and morphogenesis^25^. Besides their core of neutral lipids, LDs are made up of a phospholipid monolayer, which is decorated by various proteins involved in their biology^26–28^. In the context of pluripotency, we uncovered that the LD-associated protein CIDEA (Cell death-inducing DNA fragmentation factor alpha-like effector A) is strictly required to optimise lipid storage *in vivo* as well as in ESCs, which share the same ability as the blastocyst to form and enlarge LDs. Critically, depleting CIDEA expression leads to a loss of lipid storage and abnormal morphogenesis of pluripotent progenitors both *in vitro* and *in vivo*^25^. While the accumulation of stored lipids is recognised as an integral part of peri-implantation development, the mechanisms involved in the initial build-up of LDs and how metabolic reprogramming events integrate with other regulatory pathways remain to be investigated.

In this study, we took advantage of the inherent heterogeneity of ESCs in FBS/LIF^11,29,30^ to isolate clonal populations that harbour different amounts of LD-stored lipids. We show that low and high-LD containing ESC clones segregate along a naïve-to-primed pluripotency trajectory, validating that LD accumulation coincides with the earliest steps of differentiation as previously established in ESC-derived 3D spheroids and *in vivo*^25^. Using these clonal model systems, we demonstrate that the glycolytic switch at the exit of naïve pluripotency^13^ is preceded by an acute enhancement of oxidation coupled with *de novo* fatty acid synthesis, fuelling the lipid storage capacity of cells. Unexpectedly, we identified that ZSCAN4 (ZINC finger and SCAN domain containing 4), a key pluripotency-associated regulator of genome integrity, is preferentially active in high-LD containing clones, corroborating our observations of global DNA demethylation and heterochromatin re-organisation in these cells as indicators of ZSCAN4 events^31–34^. Mechanistically, we established that ZSCAN4 induction is downstream of CIDEA-promoted lipid storage and furthermore, that it takes place in cells transiently experiencing telomere attrition at the time of maximum LD accumulation. Importantly, inhibiting the ability of cells to accumulate LDs by CIDEA depletion abolishes both *Zscan4* induction and the shortening/re-elongation of telomeres, demonstrating that these events are co-occurring under metabolic control. Collectively, our findings uncover a metabolism-induced telomere checkpoint in ESCs transitioning from naïve-to-primed pluripotency. We propose a novel mechanism by which the genome integrity of pluripotent progenitors can be preserved as they undergo metabolic remodelling required for subsequent development.

## Results

### Lipid storage heterogeneity is a hallmark of ESCs cultured in FBS/LIF

ESCs in FBS/LIF co-exist in distinct pluripotency states from naïve and self-renewing to more primed towards differentiation, as reflected in heterogeneous transcriptional and epigenetic states^11,29,30^. We identified that ESC heterogeneity also manifests at the metabolic level with evidence of cells accumulating varying amounts of stored lipids in the form of enlarged LDs (Figure S1A,B). To address whether this delineates discrete sub-populations in FBS/LIF, we isolated and successfully established a panel of sixteen ESC clones that, under identical culture conditions, harbour different amounts and morphologies of LDs as identified by phase-contrast microscopy (Figure 1A). The co-existence of ESCs with low and high LD-stored lipid content (hereafter referred as to L-LD and H-LD respectively) was further validated by quantifying the levels of neutral lipids via BODIPY 493/503 staining (Figure 1B,C) and triglycerides (TAGs) (Figure 1D) in representative clones using confocal imaging and Liquid Chromatography Mass Spectrometry (LC-MS), respectively. While L-LD clones showed uniformly low BODIPY and TAG levels, we detected higher and more broadly distributed values amongst H-LD clones, reflecting the dynamic process of lipid accumulation.

**Figure 1:**
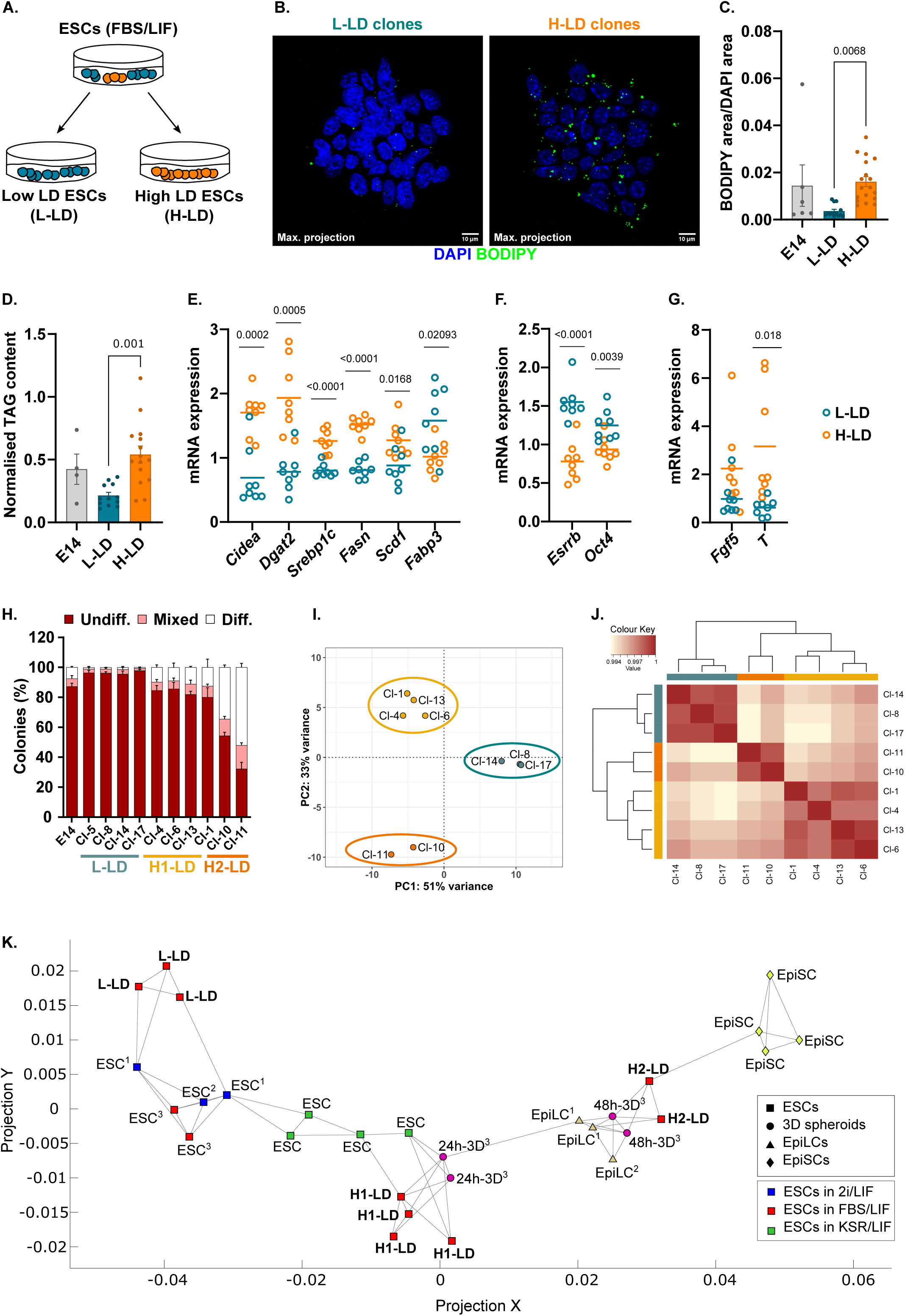
Lipid droplet heterogeneity delineates discrete pluripotency states in ESCs. (A) Isolation of low and high-containing lipid droplet (LD) ESC clones, hereafter referred as to L-LD and H-LD clones, from bulk FBS/LIF culture of E14-ESCs. (B) Representative images of LD clones in FBS/LIF stained with BODIPY 493/503-stained LDs (green) and DAPI-stained nuclei (blue). Images shown are z-stack maximum projections, scale bars, 10 μm. (C) Quantification of BODIPY signal, normalized to DAPI (area), in L-LD (8, 14, 17), H-LD (4, 6, 10, 11) clones and parental culture (E14). For each culture, n>5 colonies were imaged in independent experiments (n=3). (D) LC-MS quantification of TAG normalised to total DNA content in L-LD (8, 14, 17) and H-LD (4, 6, 10, 11) clones (n=4). Error bars, means ± s.e.m.; ANOVA with Tukey’s post hoc test. (E-G) Gene expression profiling (RT-qPCR) in LD ESC clones (passage 1-2) of genes associated with (E) lipid metabolism, (F) pluripotency, and (G) differentiation. Data shown are normalised to housekeeping genes (*S17, L19*). Each circle represents a single clone; lines represent mean expression; two-tailed t-test with Welch’s correction. (H) Percentage of colonies formed from cells seeded at low density and grown for 7 days in FBS/LIF. Colonies were counted and scored as undifferentiated, mixed, and differentiated based on alkaline phosphatase staining. Error bars, means ± s.e.m. (n=3). (I) PCA was performed on RNA-seq data collected from LD clones. (J) Correlation heatmap of RNA-seq samples showing three distinct clusters as highlighted in (I). (K) Pseudo-differentiation trajectory using dimensionality reduction of transcriptomic data from ESCs (square) in 2i/LIF (blue), FBS/LIF (red), and KSR/LIF (green); ESC-derived 3D spheroids collected at different timepoints of differentiation (circles) and epiblast-like cells (EpiLC; triangles); EpiSCs (diamond). Superscript indicates ESCs prior to and upon differentiation within the same study.

LD storage capacity is optimized in ESCs when lipids are in excess via the action of the LD-associated protein CIDEA, which mediates the fusion and hence enlargement of pre-existing LDs^25,28^. To correlate the expression of this factor with the varying amounts of neutral lipids we observed in ESCs, *Cidea* transcript levels were quantified in LD clones by RT-qPCR. Results showed a high degree of transcriptional heterogeneity across the LD clones, with the highest *Cidea* expression detected in H-LD clones (Figure 1E). Differential gene expression was also evident amongst clones when examining additional metabolic genes including *Dgat2* (diacylglycerol O-acyltransferase 2) and *Fabp3* (fatty acid binding protein 3) with roles in lipid storage and intracellular transport, respectively. These transcriptional differences were maintained upon serial passages of L-LD and H-LD clones (Figure S1C), confirming the stability of the clone phenotypes. Moreover, lowered expression of the pluripotency regulators *Esrrb* (estrogen related receptor beta) and *Oct4* (octamer-binding transcription factor 4) was noted in H-LD clones, concomitant with an increase in the induction of early differentiation markers such as *Fgf5* (fibroblast growth factor 5) and *T/Brachyury* (T-box transcription factor) (Figure 1F,G). These results point to functional differences between L-LD and H-LD clones. In agreement, we found that H-LD clones in FBS/LIF display higher incidences of spontaneous differentiation, with two sub-groups of H-LD clones forming over 10% (H1-LD) and 40% (H2-LD) of fully differentiated colonies, respectively (Figure 1H). Under “ground state” (2i/LIF) culture conditions^35^, however, H-LD clones resumed a “L-LD” like phenotype showing robust self-renewal ability together with the rare detection of small LDs as seen in routine 2i/LIF ESC culture (Figure S1A,B,D). Collectively, our findings uncover a high degree of lipid metabolic heterogeneity in FBS/LIF ESC culture, potentially underlying the priming of cells towards differentiation.

### Low and high-LD containing clones segregate along a differentiation trajectory

To further interrogate a link between LDs and pluripotency states in ESCs, we compared the transcriptomes of representative L-LD and H-LD clones using RNA-sequencing (RNA-seq). Principal component (PC) and correlation heatmap analyses of the data revealed three distinct clusters of LD clones (Figure 1I,J). As shown in Figure 1I, PC-1 separated L-LD and H-LD clones into two main clusters showing the majority of transcriptomic variance between all samples (PC-1 = 51%). PC-2 further separated the H-LD clones into two additional clusters, which corresponded to the H1-LD and H2-LD clone sub-groups we identified based on differential self-renewal capacity (Figure 1H). Moreover, examination of pathways associated with differentially expressed genes revealed the prevalence of terms pertinent to development (e.g. “Developmental biology”) in H2-LD against either H1-LD or L-LD clones (Figure S1E). In contrast, terms associated with self-renewal and pluripotency regulators (e.g. “PluriNetWork”) were enriched in L-LD clones. Importantly, lipid metabolism and other metabolic processes were confirmed as distinguishing features of LD clone clusters (Figure S1E), further highlighting the existence of metabolic heterogeneity in ESC culture.

Having established the transcriptional profiles of L-LD, H1-LD, and H2-LD clones, we asked whether these clones capture distinct cell states at the onset of differentiation. We developed a pseudo-differentiation trajectory map using dimensionality reduction clustering^36^ onto which the gene expression profiles of our clones could be projected (Figure 1K). For this, we took advantage of published RNA-seq datasets generated from ESCs grown under different culture conditions, EpiSCs and *in vitro* ESC-induced differentiation models towards formative/primed pluripotency^25,37,38^ (see also Supplemental Methods Table 1). Notably, we included the transcriptional profiles of differentiating 3D spheroids; a model system we have previously exploited to uncover the importance of lipid storage in the cellular remodelling of pluripotent progenitors at peri-implantation^25^ (Figure S1F). We confirmed that L-LD clones most closely aligned with the naïve/ground state pluripotency of ESCs in 2i/LIF (Figure 1K). H1-LD clones were revealed as an early transitional state, coinciding with maximum LD accumulation in 3D spheroids (24 hours post induction; Figure S1F,G) and ESCs grown under lipid-rich conditions known to promote LD enlargement^25^ and developmental progression^38,39^. The H2-LD clones were the furthest along the differentiation trajectory, clustering with formative ESC-induced Epiblast-like cells (EpiLC) and 48-hour 3D spheroid samples, yet not reaching the fully established primed state of EpiSCs (Figure 1K). Altogether, our results demonstrate that L-LD, H1-LD, and H2-LD clones in FBS/LIF represent discrete pluripotent and metabolic states, recapitulating the earliest steps of exit from naïve pluripotency without the requirement of any additional external stimuli.

### Enhanced oxidation signals the onset of metabolic rewiring in ESCs

To characterize the metabolic states of our clones in depth, we assessed the main metabolic fluxes running in L-LD and H-LD clones when growing at similar rates under identical culture conditions (Figure S2A). We used Seahorse assays to monitor real time rates of oxygen consumption (OCR) and extracellular acidification (ECAR) as readouts of mitochondrial respiration and glycolytic capacity, respectively (Figure 2A,B). H-LD clones exhibited higher OCR values across all mitochondrial respiration-linked parameters examined (Figure 2A; Figure S2B). Notably, the oxidative status of H1-LD clones was acutely enhanced relative to L-LD clones as also confirmed by measuring total reactive-oxygen species (ROS) levels (Figure 2C). Concurrently, the glycolytic capacities of H-LD clones were accelerated, especially of H2-LD clones (Figure 2B; Figure S2C), in agreement with the more developmentally advanced states of these cells^13^. This was also demonstrated by increased glutamine dependence for mitochondrial respiration in H-LD clones alone (Figure S2D,E), an additional hallmark of cells transitioning toward primed pluripotency^40,41^. These results delineate transitional metabolic phenotypes in between ESCs and EpiSCs at the exit of naïve pluripotency.

**Figure 2:**
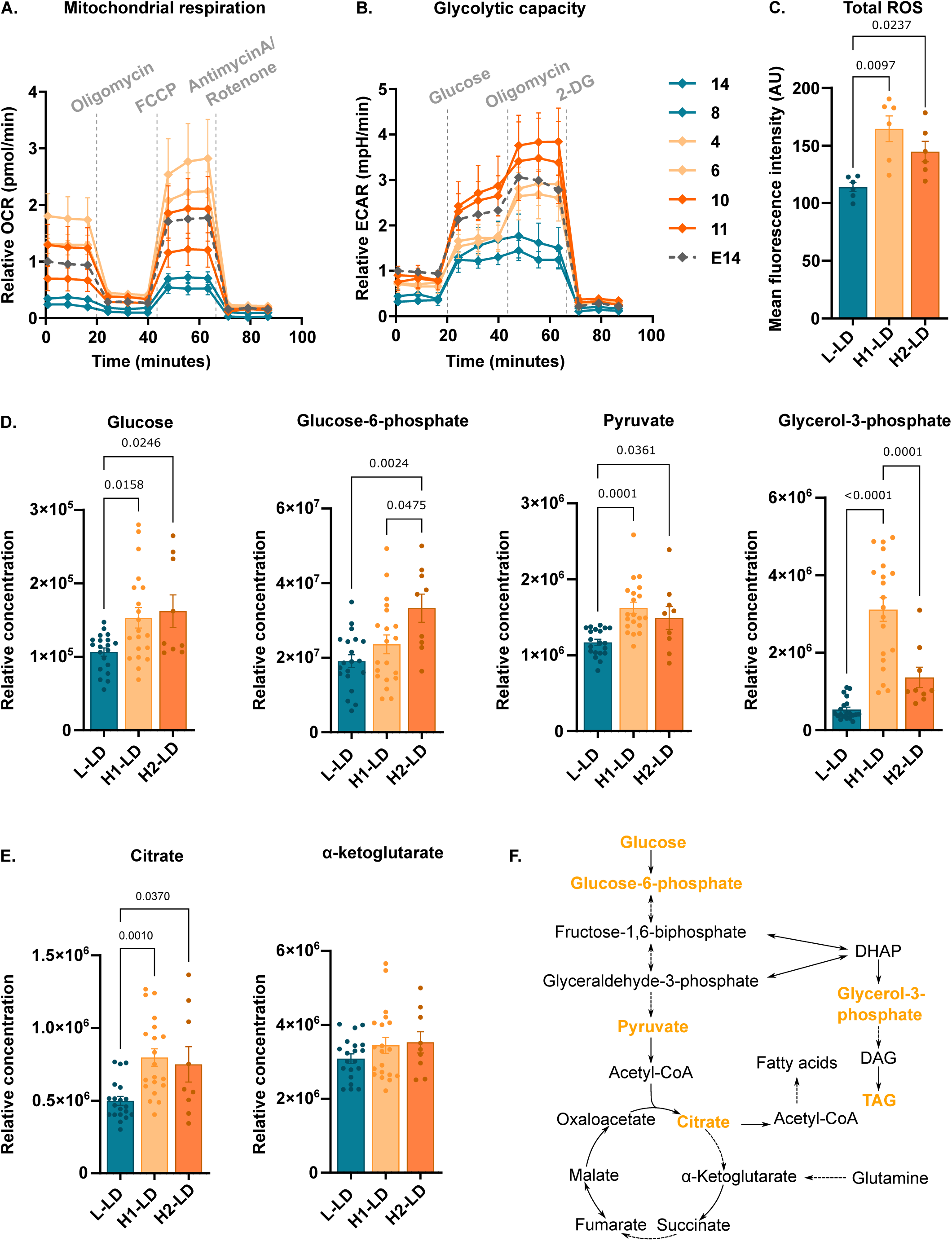
Enhanced oxidation signals the onset of metabolic rewiring in ESCs. Seahorse analysis of (A) OCR and (B) ECAR as measures of mitochondrial respiration and glycolytic capacity in specified LD clones. Data are normalised to parental line (E14; dashed line). Indicated drugs were added at time points shown. Error bars, means ± s.e.m. (n=3). (C) Total ROS levels, indicated by DCFDA fluorescence, in L-LD (8), H1-LD (4), and H2-LD (11) clones using flow cytometry; Error bars, means ± s.e.m.; ANOVA with Tukey’s post-hoc test (n=6). (D, E) Relative concentration of (D) glycolysis and (E) TCA cycle intermediate metabolites in L-LD (5, 8, 14, 17), H1-LD (1, 4, 6, 13) and H2-LD (10, 11) measured by GC-MS. Error bars, means ± s.e.m.; ANOVA with Tukey’s post-hoc test (n=5 per clone). (F) Schematic of metabolic pathways. Dashed arrows indicate intermediate steps excluded.

To complement our Seahorse experiments, we performed untargeted Gas Chromatography Mass Spectrometry (GC-MS) metabolomics. Supporting our earlier data, dynamic changes in the redox status of H-LD clones were indicated by the activation of antioxidant responses and hence lower detection of reduced methionine and glutathione metabolites in these cells (Figure S2F,G). Moreover, we observed a significant increase in glucose uptake and concentration of glycolytic intermediates in H-LD compared to L-LD clones, in agreement with glycolysis acceleration (Figure 2D,F). Notably, higher concentrations of the TAG-precursor, glycerol-3-phosphate (G3P), were detected in H1-LD and to a lesser extent in H2-LD clones, mirroring the levels of TAGs detected in these ESC clones (Figure 1D and Figure S2H,I for direct comparison). Given the enrichment of pathways associated with the citric acid (TCA) cycle in H-LD clones (Figure S1E), we also surveyed the representation of TCA intermediate metabolites. We identified a selective and significant increase in the concentration of citrate in H-LD clones in contrast to α-ketoglutarate shown in Figure 2E. This suggests a shift in the engagement of the TCA cycle toward citrate production in more developmentally advanced clonal populations. Taken together, our complementary analyses provide evidence for an acceleration in both mitochondrial respiration and citrate production in ESCs at the exit of naïve pluripotency as they transition towards a more glycolytic phenotype.

### LD accumulation reflects higher TCA-fuelled lipid synthesis in ESCs

Given the lipid-rich phenotype of H-LD clones, we asked if the higher production of G3P and citrate we observed in these clones was linked to lipid biosynthetic processes. Using the transcriptome data from LD clones, we surveyed the expression of key genes involved in *de novo* lipogenesis (Figure 3A,B). These included *Acly*, which encodes the citrate ATP lyase (ACLY) enzyme that links the TCA cycle to fatty acid (FA) synthesis and provides the main source of acetyl-CoA required for lipid biogenesis. Additionally, we examined the expression of *Acaca* (acetyl-CoA carboxylase) and *Fasn* (fatty acid synthase), encoding for two rate limiting enzymes in the synthesis of FAs, *Scd1* (stearoyl-CoA desaturase 1) involved in the formation of monounsaturated FAs essential for ESC survival^42^, and *Srebf1*, which encodes the sterol regulatory element-binding protein SREBP1 that activates the expression of *Acly*, *Fasn* and *Scd1*. All these genes were consistently upregulated in H1-LD and H2-LD relative to L-LD clones as seen for *Dgat2* involved in the formation of TAGs (Figure 3A,B). Enhanced activity of SREBP1 was also confirmed at the protein level, showing increased ratios of SREBP1 cleaved versus precursor proteins in H-LD clones (Figure 3C; Figure S2J for quantification). This, together with higher FASN protein expression (Figure 3C; Figure S2K), is suggestive of a *de novo* lipid synthesis switch in H-LD clones.

**Figure 3:**
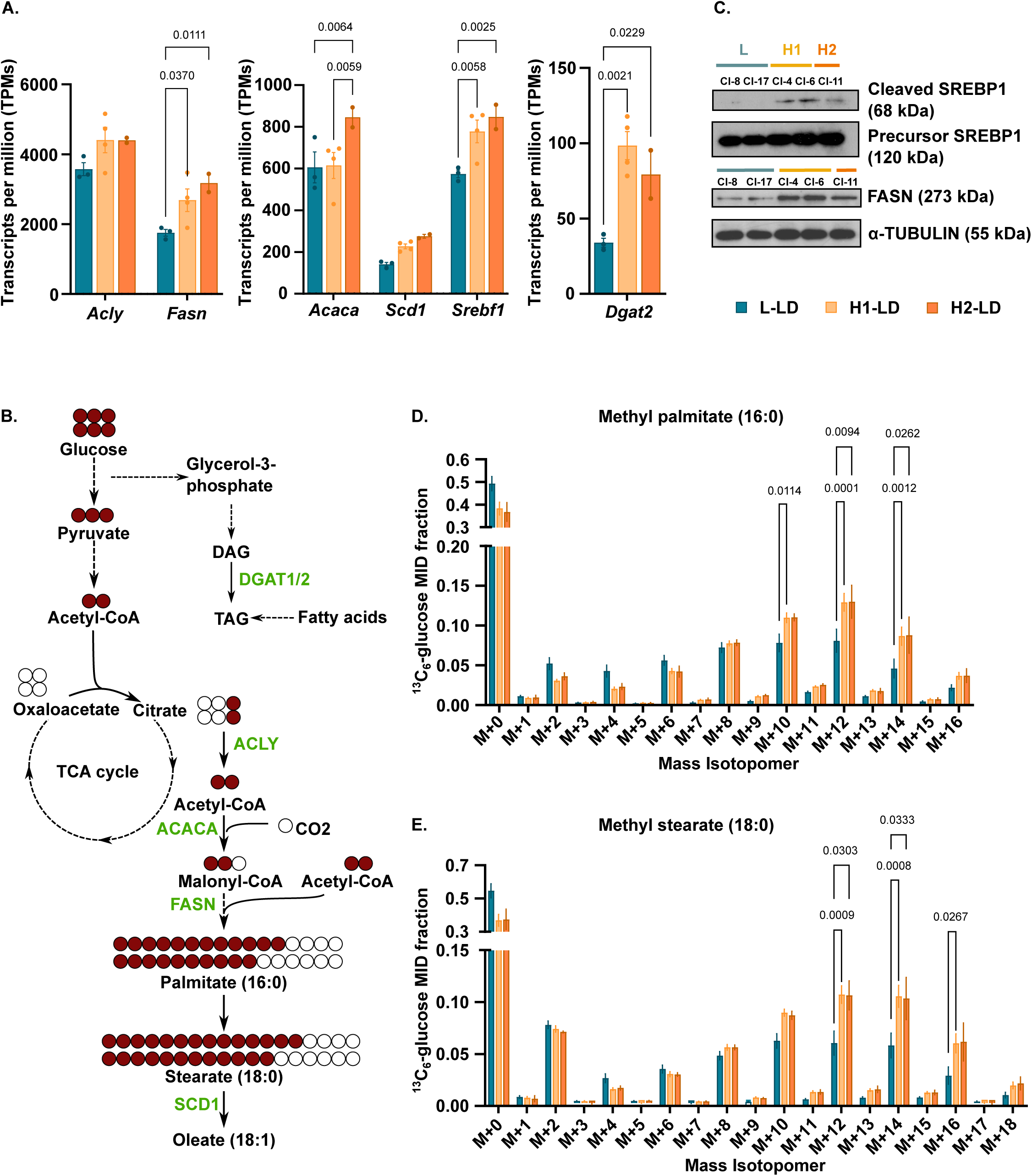
LD accumulation reflects higher TCA-fuelled lipid synthesis in ESCs. (A) Gene expression profiling (RNA-seq) in L-LD (8, 14, 17) and H-LD (1, 4, 6,10, 11, 13) clones of genes associated with fatty acid synthesis and storage, represented as transcript per million (TPMs). Error bars, means ± s.e.m.; Fisher’s two-way ANOVA test. (B) Schematic showing the incorporation of ^13^C_6_-glucose-drived carbon (red) in fatty acid synthesis and elongation. Key enzymes are shown in green; dotted arrows indicate intermediate steps excluded. (C) Representative western blot of cleaved and precursor SREBP1, FASN, and α-TUBULIN used as loading control (see also Figure S2J,K). (D,E) ^13^C_6_-labelled FAME represented as a mass isotopomer distribution (MID) of (D) methyl palmitate and (E) methyl stearate in LD clones. Error bars, means ± s.e.m.; ANOVA with Tukey’s post-hoc test (n=3 per clone).

FA metabolism can also be monitored by tracing ^13^C_6_-glucose, which generates acetyl-CoA (from cytosolic citrate) containing two labelled carbons to be incorporated into FAs during *de novo* synthesis (producing palmitate) and elongation (producing stearate) (Figure 3B). Analysis of total FAs detected (labelled and unlabelled) by GC-MS showed no significant difference in the concentrations of methyl palmitate (16:0) and methyl oleate (18:1) across the three clonal populations (Figure S2L). However, we noted an increase in the levels of methyl stearate (18:0) in H1-LD clones. Stearate distinguishes naïve from formative/primed pluripotency^43^, reflecting the different functional states of our clones. To directly test the lipogenic activity of the clones, we determined the utilisation of ^13^C_6_-glucose for the biosynthesis of methyl palmitate and methyl stearate via mass isotopomer distribution (MID) analysis (Figure 3D,E). Results showed enhanced rates of *de novo* FA synthesis in both H1-LD and H2-LD clones, as indicated by the lower fraction of unlabelled species (M+0). Concomitantly, H-LD clones showed significantly higher enrichment of ^13^C_6_-glucose in methyl palmitate (M+10, M+12, M+14) and methyl stearate (M+12, M+14, M+16), suggesting increased utilisation of glucose-derived acetyl-CoA for FA synthesis and elongation in these clones. Collectively, these findings demonstrate a shift in TCA cycle engagement towards citrate production and *de novo* lipid synthesis at the exit of naïve pluripotency, most likely facilitating the accumulation of LDs required for subsequent differentiation.

### ChEP-based profiling of chromatin-bound proteomes in LD clones

One notable feature of L-LD and H-LD clones is the stability of their phenotypes upon propagation (Figure S1C), suggesting they maintain different epigenetic programmes. To gain further insights, we profiled the landscapes of chromatin-bound proteins in representative clones by Chromatin Enrichment for Proteomics (ChEP)^44^. Data was processed to generate label-free quantification (LFQ) values and filtered for potential contaminants and lowly expressed peptides. As an important validation, we confirmed that L-LD and H-LD clones could be segregated based on the data using PCA analysis (Figure S3A). Among differentially bound epigenetic regulators, we observed an enrichment of DNA methylation-associated proteins including DNMT1 (DNA methyltransferase 1) and UHRF1 (ubiquitin-like, containing PHD and RING finger domains 1) in L-LD clones compared to either H1-LD or H2-LD clones (Figure 4A,B). Most remarkably, this was reflected in H-LD clones by a converse enrichment of ZSCAN4 protein, which is directly implicated in the degradation of DNMT1 and UHRF1 in ESCs^34^. Accordingly, we confirmed by western blotting that H-LD clones express higher levels of ZSCAN4 and reduced levels of DNMT1 and UHRF1 compared to L-LD clones (Figure 4C). This corroborates with lower levels of global DNA methylation in H-LD clones as quantified by High-Performance Liquid Chromatography-Mass Spectrometry (HPLC-MS) (Figure 4D).

**Figure 4:**
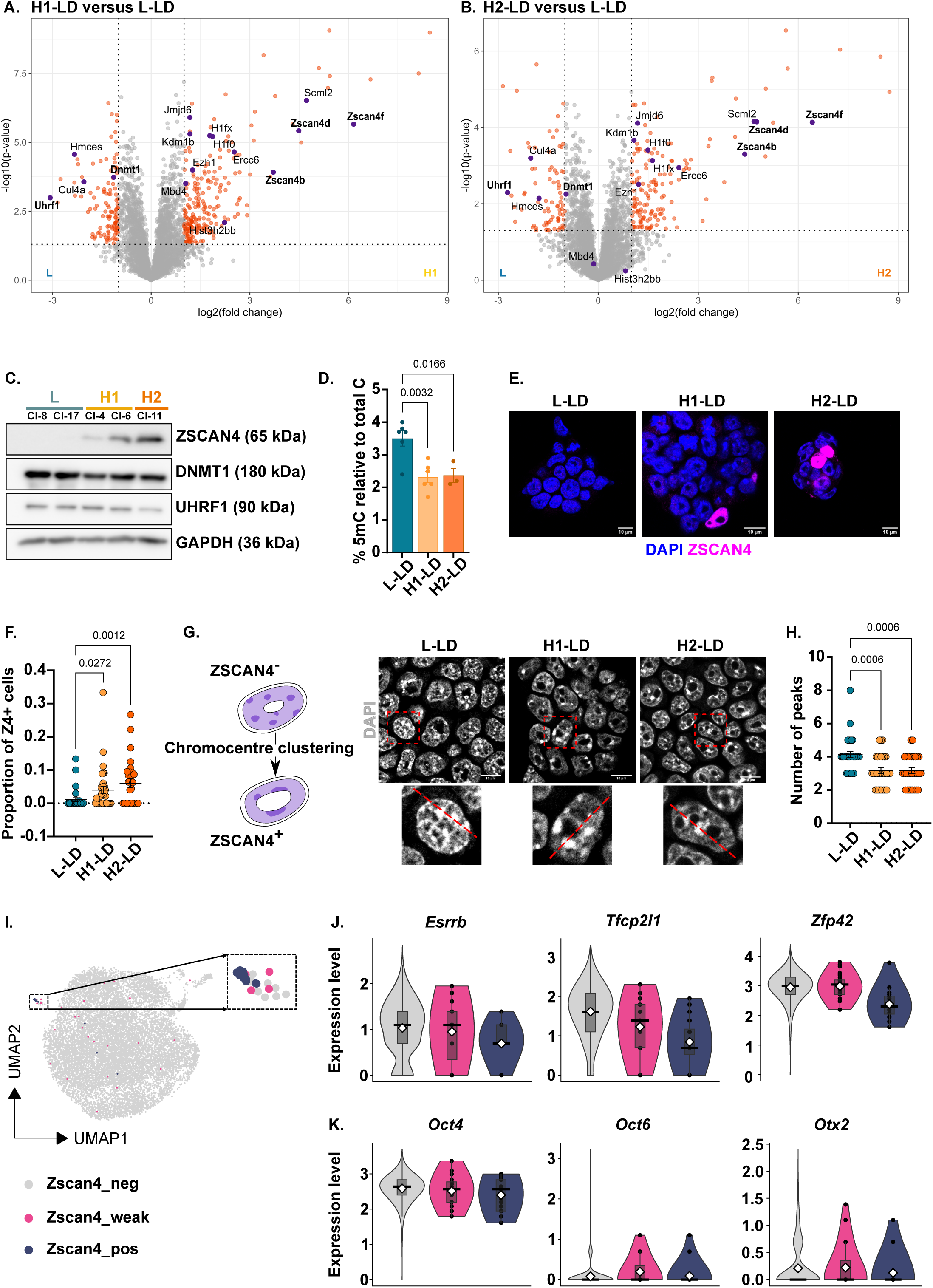
Lipid storage and *Zscan4* induction intersect at the onset of differentiation. (A,B) Differential enrichment of chromatin-bound proteins in L-LD (8, 17), H1-LD (4,6) and H2-LD (11) clones as determined by ChEP. Data points shown in red indicate p-value < 0.05, log2 fold change ± 1 cut-off. (C) Representative western blot of LD clones for ZSCAN4, DNMT1, UHRF1 and GAPDH used as loading control (n=3). (D) Percentage of 5-methylcytosine (5mC) to total cytosine (C) in L-LD (8, 17), H1-LD (4, 6) and H2-LD (11) clones as quantified by HPLC-MS. Error bars, means ± s.e.m.; ANOVA with Tukey’s post-hoc test (n=3 per clone). (E) Representative immunofluorescence images for ZSCAN4 and DAPI in L-LD (8), H1-LD (4) and H2-LD (11) clones; scale bars, 10 μm. (F) Proportion of ZSCAN4-positive (Z4+) to total number of cells per colony as shown in (E). Each dot represents one colony; Error bars, mean ± s.e.m.; ANOVA with Tukey’s post-hoc (n>30 colonies across 3 independent experiments). (G) Schematic of differential chromocenter clustering in ZSCAN4-negative (−) and ZSCAN4-positive (+) ESCs (left) and intensity linescan analysis of chromocenters stained with DAPI in LD clones as indicated in insets (right); scale bars, 10 μm. (H) Quantification of chromocenters numbered as peaks in intensity linescan profiles. Each dot represents one nucleus; lines indicate mean ± s.e.m.; ANOVA with Tukey’s post-hoc test (n=30 nuclei across 3 independent experiments). (I) Single-cell RNA-seq and UMAP analysis showing identification of *Zscan4*-negative (grey), weakly-expressing (pink), and positive (purple) ESCs across L-LD (8, 17), H1-LD (4, 6) and H2-LD (11) clones and parental ESC (E14) culture. (J,K) Expression level of naïve (*Esrrb, Tfc2pl1, Zfp42*), core (*Oct4*), and formative/primed (*Oct6, Otx2*) pluripotency genes in *Zscan4*-negative, weakly expressing, and positive single cells shown as violin plots.

Originally identified as a 2-cell (2C) mouse embryo marker^31^, ZSCAN4 is also sporadically expressed in ESC culture, playing key roles in the maintenance of genome integrity and telomere homeostasis^32,33,45–48^. Accordingly, ZSCAN4 protein was detected only in a fraction of cells in all clonal populations (Figure 4E). However, the frequency of these ‘ZSCAN4 events’ was increased in H1-LD and most significantly in H2-LD clones compared to their L-LD counterparts (Figure 4F). This trend was also reflected at the transcriptional level with the upregulation of *Zscan4* paralogues in H-LD clones, together with *Dppa3* (developmental pluripotency associated 3) (Figure S3B,C), implicated in DNA demethylation in ZSCAN4-expressing ESCs^49^. In comparison, *Dppa2*, *Dppa4*, and the 2C embryonic marker *Duxf3* (double homeobox)^50,51^ were not upregulated in H-LD clones. In addition to DNA demethylation, ZSCAN4 events coincide with changes in heterochromatin organisation^33,34^. Therefore, we monitored the number and distribution of chromocenters (heterochromatin foci) in L-LD and H-LD clones (Figure 4G). Using DAPI linescan analysis^52,53^, we observed a decrease in the number of chromocenters in H-LD clones, forming larger foci positioned around nucleoli (Figure 4G,H), indicative of chromatin decompaction^33^. Taken together, these analyses reveal a higher incidence of ZSCAN4 events in lipid-rich H-LD clones, transiently promoting DNA demethylation and chromatin accessibility^33,34^. We propose that these global epigenetic changes may at least partly account for the stability of our clone phenotypes.

### Lipid storage and *Zscan4* induction in ESCs intersect at the single-cell level

Unexpectedly, our results indicate that ZSCAN4 is expressed more frequently in ESCs undergoing metabolic and cell state transitions towards primed pluripotency. To further understand the relationship between *Zscan4* induction and developmental progression, we applied single-cell transcriptomics. Representative LD clones along with parental E14-ESCs were processed using the 10X Genomics technology. After quality control and filtering of the data, we retrieved between 2,465 and 3,796 single-cell transcriptomes from LD clone and parental cultures. To verify that the data recapitulated the key phenotypic features of L-LD, H1-LD and H2-LD clones, we computed “naïve”, “primed” and “lipid” signature scores based on genes associated with these features, confirming higher “primed” and “lipid” scores in H1-LD and H2-LD cells (Figure S3D-F). In all populations examined, *Zscan4* transcripts were detected only in a few cells (n= 4-27), as previously reported in single-cell datasets collected from early differentiating ESCs^54^. However, differences in total *Zscan4* UMI counts could be discerned amongst our samples, being consistently lower in L-LD cells as anticipated (Figure S3G).

Subsequent UMAP analysis of merged samples highlighted a separate cell cluster consisting of *Zscan4*-positive (purple) and *Zscan4*-negative (grey) cells, and cells expressing weaker levels of *Zscan4* transcripts (pink) (Figure 4I). Inferring a link with ZSCAN4 events, genes typically upregulated in ZSCAN4-expressing ESCs^48,49,51,55,56^ were enriched in this cluster and all individual *Zscan4*-positive cells detected in our analysis (Figure S3H). Interestingly, repressors of ZSCAN4 events such as *Chaf1a* (chromatin assembly factor 1 subunit A)^57^ and *Rif1* (replication timing regulatory factor 1)^58^ were more strongly expressed in *Zscan4*-weak and *Zscan4*-negative cells, reflecting the tight control of *Zscan4* induction. When examining the pluripotency status of these cells, we found that *Zscan4*-positive cells express the lowest levels of naïve pluripotency genes (Figure 4J). However, they retain relatively high expression of *Oct4* together with the incidental induction of the formative markers *Oct6* (octamer-binding transcription factor 6) and *Otx2* (orthodenticle homeobox 2) (Figure 4K), indicative of an early transitional signature^59,60^. Importantly, we identified that *Zscan4*-expressing cells exhibit the highest “lipid” scores in H-LD populations (Figure S3I), where they are also more frequently detected and display higher *Zscan4* transcript levels (see Figure S3G). This reinforces the proposed association between lipid storage and ZSCAN4 events at the single-cell level. Collectively, these data reiterate that, while *Zscan4* induction is enhanced in lipid-rich cells at the onset of differentiation, it remains a dynamic and tightly regulated event.

### ZSCAN4 induction is driven by anabolic activity in ESCs and *in vivo*

To directly test if the ZSCAN4 phenotype is stimulated by the lipid-rich state of ESCs, we made use of a previously established cell line that ectopically expresses *Cidea* to enforce lipid storage^25^ (Figure S4A). *Cidea* overexpressing (CIDEA OE) ESCs shared several characteristics with H1-LD clones, including an accumulation of enlarged LDs (Figure 5A,B), enhanced oxygen consumption, and an upregulation of lipid biosynthesis markers (pSer455-ACLY, FASN) (Figure S4B-D). Most importantly, we demonstrated that enforcing lipid storage in ESCs was sufficient to increase the frequency of ZSCAN4 events as evidenced by a higher proportion of ZSCAN4-positive (Z4+) cells and global DNA demethylation in CIDEA OE compared to control cultures (Figure 5C,D; Figure S4E). In contrast, ectopically expressing *Zscan4c* itself neither promoted mitochondrial respiration nor enforced a lipid-rich state in ESCs. While *Zscan4c* overexpression (ZSCAN4 OE) significantly reduced the protein expression and activity of DNMT1 and UHRF1 in ESCs, it had no impact on the amount and morphology of LDs and metabolic fluxes running in these cells (Figure 5E-H; Figure S4F-I). Enhanced *Zscan4* expression, however, was associated with early changes in the pluripotency status of cells as denoted by *Otx2* upregulation in both CIDEA and ZSCAN4 OE ESCs (Figure 5I). Taken together, these experiments demonstrate that ZSCAN4 induction is downstream of a lipid biosynthesis switch in ESCs.

**Figure 5:**
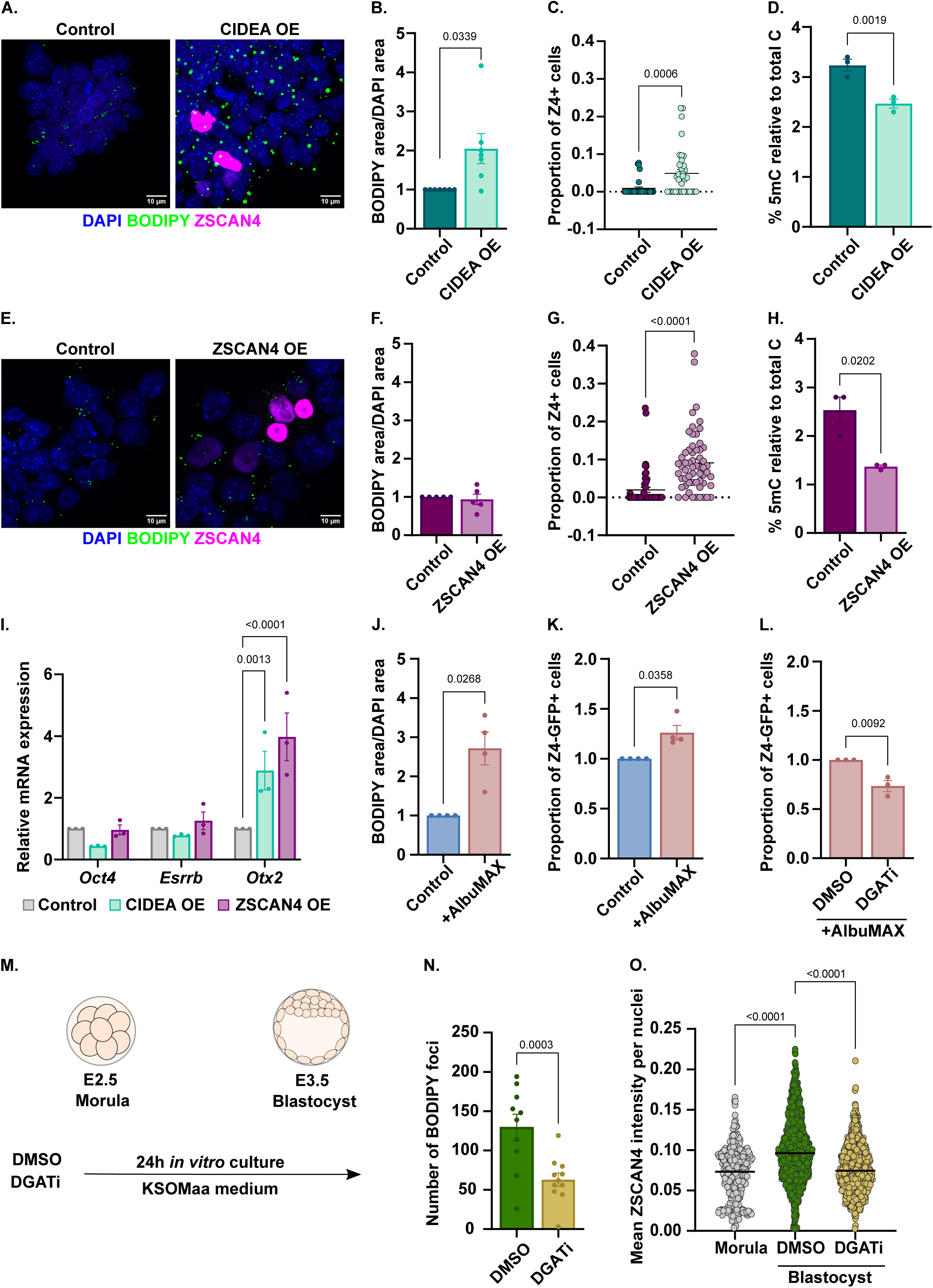
ZSCAN4 induction is driven by anabolic activity in ESCs and *in vivo*. (A) Representative immunofluorescence images for ZSCAN4, BODIPY and DAPI in *Cidea* overexpressing (CIDEA OE) and control (empty CAG vector) ESCs; scale bars, 10 μm (B) Quantification of BODIPY signal in CIDEA OE and control cultures. Error bars, means ± s.e.m.; paired, two-tailed t-test with Welch’s correction (n=7). (C) Proportion of ZSCAN4-positive (Z4+) cells in CIDEA OE and control cultures; two-tailed t-test with Welch’s correction (n>30 colonies across 3 independent experiments). (D) Percentage of 5mC to total C in CIDEA OE and control ESCs. Error bars, means ± s.e.m.; paired two-tailed t-test with Welch’s correction (n=3). (E) Representative immunofluorescence images for ZSCAN4, BODIPY and DAPI in *Zscan4* overexpressing (ZSCAN4 OE) and control ESCs; scale bars, 10 μm. (F) Quantification of BODIPY signal in ZSCAN4 OE and control cultures. Error bars, means ± s.e.m.; two-tailed t-test with Welch’s correction (n=5). (G) Proportion of Z4+ cells in ZSCAN4 OE and control cultures; two-tailed t-test with Welch’s correction (n>30 colonies across 3 independent experiments). (H) Percentage of 5mC to total C in ZSCAN4 OE and control ESCs. Error bars, means ± s.e.m.; paired two-tailed t-test with Welch’s correction (n=3). (I) Gene expression profiling (RT-qPCR) in CIDEA OE and ZSCAN4 OE ESCs of pluripotency genes, shown relative to control ESCs. Error bars, means ± s.e.m.; Fisher’s two-way ANOVA test (n=3). (J) Quantification of BODIPY signal in E14-ESCs under control (FBS/LIF) and AlbuMAX (FBS/LIF + 1% AlbuMAX) conditions for 48 hours. Error bars, means ± s.e.m.; paired two-tailed t-test with Welch’s correction (n=4). (K,L) Proportion of ZSCAN4::GFP positive (Z4-GFP+) ESCs using flow cytometry and E14-ZE3 reporter line in (K) control and AlbuMAX conditions and (L) AlbuMAX with DMSO or inhibitors against DGAT1/2 (DGATi) for 48 hours. Error bars, means ± s.e.m.; paired two-tailed t-test with Welch’s correction (n=3-4). (M) Schematic of mouse embryo *in vitro* culture experiments. (N) Quantification of BODIPY-stained foci in control (DMSO) and DGATi (25 μM)-treated cultured blastocysts. Each dot represents one embryo. Error bars, means ± s.e.m.; unpaired two-tailed t-test with Welch’s correction (n=10-11). (O) Mean ZSCAN4 intensity per nuclei in morula, and DMSO- and DGATi-treated cultured blastocysts. Each dot represents an individual nucleus, with the median value indicated. ANOVA with Dunnett’s post-hoc test to compare to DMSO control.

To further determine the importance of lipid storage in the regulation of *Zscan4*-expressing cells, we constructed an ESC line that contains stable integration of the *Zscan4c* promoter-driven EGFP fluorescent reporter (ZE3) (Figure S4J). This reporter has been previously described to reflect the dynamic expression pattern of their endogenous counterpart^32,46,61^. While ~0.1-1.5% of ESCs were labelled by the reporter in FBS/LIF, substituting FBS by Knockout Serum Replacement (KSR) and thereby growing cells in lipid-enriched medium increased the proportion of Zscan4c::EGFP-positive (Z4-GFP+) cells over 72 hours (Figure S4K), along with the rapid induction of *Cidea* (Figure S4L) and LD accumulation^25^. A short treatment with AlbuMAX alone (i.e. the lipid component of KSR) was sufficient to achieve a similar effect (Figure 5J,K). However, this effect was reduced upon addition of inhibitors against DGAT1/2 (DGATi; Figure 5L), demonstrating a direct role for lipid storage in the regulation of ZSCAN4 events.

*In vivo*, LDs are actively enlarged and accumulated during the morula-to-blastocyst transition prior to implantation and differentiation^22–25^. We asked if the build-up of LDs also influences the expression pattern of ZSCAN4 in this context. For this, we harvested morula embryos (E2.5; ~ 74h post-hCG injection) and cultured them up to the early blastocyst stage (E3.5; ~ 98h post-hCG) with DMSO or DGATi (Figure 5M). While DGATi treatment significantly lowered BODIPY signals in a dose-dependent manner, the highest dose of inhibitors increased the incidences of arrested and/or abnormal embryos (Figure 5N; Figure S5A-C) and therefore was not used for subsequent analysis. Flushed morula and cultured blastocysts (n=10-11) were stained for ZSCAN4, F-ACTIN and DAPI followed by quantitative image analysis (Figure S5D,E). We found that nuclear ZSCAN4 fluorescence intensity significantly increased from the morula to the blastocyst stage, a pattern abrogated under DGATi treatment (Figure 5O) as seen in ESCs. Collectively, our analyses uncover a causal link between enhanced lipid storage and ZSCAN4 induction at a key transitional point of development.

### A metabolism-induced telomere checkpoint operates at the onset of differentiation

Knowing that ZSCAN4 regulates the genome integrity of ESCs^32,33,45,47^, we asked if the acceleration of anabolic outputs in lipid-rich cells augments the risk for DNA damage as a possible trigger for ZSCAN4 induction. To explore this, we determined global levels of γH2AX by western blotting and the incidence of apoptosis in CIDEA OE ESCs and found that these cells did not show increased DNA damage and/or cell death, respectively (Figure S6A,B). Interestingly, however, we found that CIDEA OE ESCs exhibited shortened telomeres as assessed using quantitative telomere FISH (Telo-FISH) on chromosome metaphase spreads (Figure 6A,B). To confirm this observation, we quantified the average lengths of telomeres using qPCR-based measurements and furthermore extended our analysis to L-LD and H-LD clones (Figure 6C; Figure S6C). While our results offer evidence that telomere attrition operates in the context of high lipid storage (CIDEA OE and H1-LD), they also indicated that telomere lengths may be regulated (H2-LD) at the onset of differentiation. This suggests that a telomere elongation mechanism could be activated as cells undergo metabolic remodelling and progress towards primed pluripotency.

**Figure 6:**
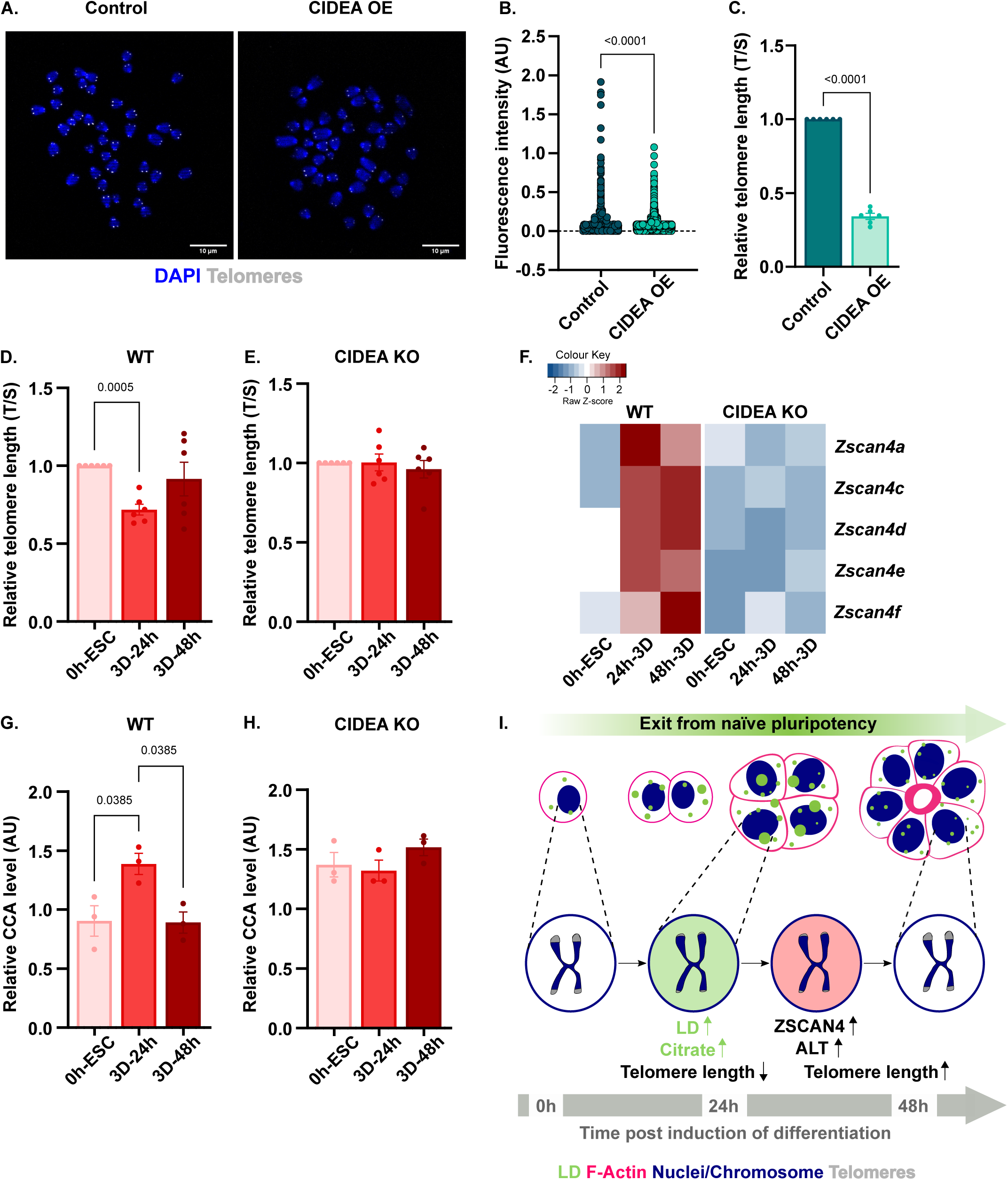
A metabolism-induced telomere checkpoint operates at the onset of differentiation. (A) Representative images of Telo-FISH using TelC-Cy3 probe and DAPI on metaphase spreads prepared from CIDEA OE and control ESCs. Scale bars, 10 μm. (B) Quantification of fluorescence intensity of Telo-FISH signal relative to DAPI signal for each telomeric foci; each circle represents one telomeric foci. n>100 across 3 independent experiments; two-tailed t-test with Welch’s correction. (C) Relative telomere length as measured by telomere qPCR, shown as a ratio of telomere to single-copy gene (T/S). Error bars, means ± s.e.m.; paired, two-tailed t-test with Welch’s correction (n=6). (D, E) Relative telomere length in undifferentiated ESCs (0h) and 3D spheroids at different timepoints of differentiation (24h, 48h) formed by (D) wild-type (WT) and (E) *Cidea* knock-out (CIDEA KO) R1-ESCs. Error bars, means ± s.e.m; ANOVA with Tukey’s post-hoc (n=6) (F) Heatmap showing relative transcript expression (RNA-seq) of *Zscan4* paralogues prior to and upon 3D spheroid formation by WT and CIDEA KO ESCs. (G,H) Relative CCA levels as measured by ALT-specific qPCR prior to and upon 3D spheroid formation by (G) WT and (H) CIDEA KO ESCs. Error bars, means ± s.e.m; ANOVA with Tukey’s post-hoc (n=3) (I) Proposed model linking metabolic rewiring and telomere length safeguarding at the exit from naïve pluripotency via the activation of ZSCAN4-associated ALT mechanisms.

To establish that telomere length is indeed actively safeguarded during the transition from naïve-to-primed pluripotency, we measured the average telomere lengths of ESCs prior to and upon induction of 3D spheroid differentiation (Figure S1F). Concomitant with maximum LD enlargement^25^, we identified a transient shortening of telomeres in 24-hour 3D spheroids formed by wild-type (WT) ESCs (Figure 6D). Telomere lengths were then restored in late differentiating cultures (48 hours), recapitulating the telomere dynamics we observed in LD clones (Figure S6C). In contrast, 3D spheroids formed by *Cidea* knock-out (CIDEA KO) ESCs, which are unable to enlarge LDs^25^, showed no shortening or significant variation in telomere lengths upon differentiation (Figure 6E). Importantly, this agrees with the loss of *Zscan4* induction in CIDEA KO 3D spheroids in contrast to WT cultures (Figure 6F). Together, these results strengthen our conclusion that lipid storage, *Zscan4* induction and telomere homeostasis are functionally connected at the onset of differentiation.

ZSCAN4 has been shown to induce telomere elongation through the Alternative Lengthening of Telomeres (ALT) in ESCs^34^, a mechanism that elongates telomeres via homologous recombination and is also active in some cancer cells^47,62,63^. To gain further understanding, we performed Telo-FISH in 3D spheroids to determine the level of telomere clustering as an indication of homologous recombination through ALT. We found a significant increase in the area of telomere foci, peaking in 24-hour 3D spheroids (Figure S6D,E) and coinciding with the onset of *Zscan4* induction (Figure 6F). The activation of the ALT mechanism was further validated in H-LD clones and 3D spheroids using two alternative assays. These included the C-circle amplification (CCA) assay, a PCR-based assay designed to detect C-circles, a hallmark of ALT activity^64^ (Figure 6G), and the formation of ALT-associated PML bodies (APBs) at telomeres^65^ detected by immunofluorescence with co-staining for PML and TRF1, a component of the telomeric Shelterin complex (Figure S6F). All these analyses concurred to demonstrate the incidence of a transient shortening of telomeres and their subsequent ALT-mediated elongation at the exit of naïve pluripotency as seen in 2D (Figure S6C,F,G) and 3D cultures (Figure 6G; Figure S6D,E,H) unless CIDEA-promoted lipid storage was abrogated (Figure 6H). Altogether, our findings reveal for the first time that telomere length is actively modulated in pluripotent progenitors at the onset of differentiation and that the activation of a telomere safeguarding mechanism via ZSCAN4 can mitigate the impact of key metabolic remodelling events taking place during peri-implantation development.

## Discussion

The transition from naïve-to-primed pluripotency is associated with extensive cellular and biochemical remodelling^5,13,25,30,38^. In this study, we identify novel metabolic states that underline early transitional steps towards the acquisition of primed pluripotency and the associated glycolytic switch. We show that high-LD containing ESCs have increased metabolic activity and capacity. Transitioning out of naïve pluripotency, these cells exhibit an acceleration of mitochondrial respiration and citrate production, fuelling FA synthesis and storage. Moreover, we find that the abilities to actively synthesise and store lipids is acquired by a subset of ESCs maintained under identical culture conditions, indicative of intrinsic metabolic reprogramming. These findings are supported by a recent study showing that the inactivation of ACLY and FASN enzymes can block the exit of naïve pluripotency in ESCs^66^. *De novo* lipid switches have also been reported as a prerequisite for adult progenitor function in neurogenesis and myogenesis^67,68^, further highlighting the fundamental importance of lipid metabolism during embryonic and postnatal development.

Unexpectedly, we uncover that the build-up of LDs that operates during pluripotency maturation is coupled with the activity of ZSCAN4 and telomere safeguarding. We find that ZSCAN4 events are promoted in the context of anabolic hyperactivity, but not from increased DNA damage and/or cell death. Mechanistically, we demonstrate that enforcing lipid storage in ESCs via *Cidea* overexpression increases the frequency of ZSCAN4 induction, leading to DNA demethylation and heterochromatin decompaction. However, these events remain dynamic and transient, suggesting that they may be confined to cells experiencing higher levels of stress^32,61^. Accordingly, we establish that H-LD clones and 3D spheroids transiently exhibit shortened telomeres at the exit of naïve pluripotency, which concurs with maximum LD accumulation, the induction of *Zscan4,* and subsequent activation of the ALT-mechanism, as also correlated at the single-cell level (see Figure S3G-I). Importantly, inhibiting lipid storage alone is sufficient to abolish telomere attrition and ZSCAN4 induction, in agreement with these events co-occurring under metabolic control. Collectively, our findings reveal a previously unrecognized mechanism by which LD accumulation, required for subsequent development^25^, is tightly co-ordinated with a metabolism-induced telomere checkpoint to preserve the genome integrity of pluripotent progenitors at the onset of differentiation (see proposed model in Figure 6I).

*In vitro*, ZSCAN4-associated telomere lengthening has been observed in steady state ESCs^32,34,45,47^ and upon somatic cell reprogramming towards naïve pluripotency^69,70^. A similar role for ZSCAN4 during the naïve-to-primed pluripotency transition has thus far not been reported. *In vivo*, ZSCAN4 is transiently expressed in mouse 2C-stage embryos where it is involved in zygotic genome activation^31^. The contribution of ZSCAN4 in regulating the rapid telomere lengthening of cleavage-stage embryos remains to be clarified. We show that ZSCAN4 can also be detected during the morula-to-blastocyst transition, which experiences a substantial accumulation of enlarged LDs^22,23,25^. Interrupting the build-up of LDs during this transition resulted in reduced levels of ZSCAN4 detection, indicating that ZSCAN4 induction may also be under metabolic control in the cavitating blastocyst as seen in ESCs. Previous reports have suggested that the blastocyst actively elongates its telomeres, potentially through ALT and telomerase reverse transcriptase (TERT)-mediated pathways^71,72^. Moreover, the depletion of *Dcaf11*, a positive regulator of *Zscan4*, significantly reduces the telomere lengths of KO blastocyst embryos *in vivo*^72^, implying a role for ZSCAN4 in telomere elongation at this developmental stage. Together, these findings along with our study using ESC-based models demonstrate that telomere length is actively modulated at the onset of implantation and differentiation. These findings also reinforce the view that ZSCAN4 reactivation may play a prime role during cell state transitions even after the 2C-embryonic stage.

An intriguing question arising from our study is how telomere length is transiently shortened in the context of accelerated mitochondrial respiration and anabolic outputs in pluripotent progenitors. Mounting evidence suggests a close connection between mitochondrial metabolism and telomere homeostasis in development and diseases^73–77^. In human ESCs, shorter telomeres have been correlated at the single-cell level with higher expression of key regulators of oxidative metabolism and lower expression of factors known to stabilize the recruitment of telomerase and shelterin complexes to telomeres^78^. The TERT protein itself has been proposed to accumulate into mitochondria rather than the nucleus when needed to protect cells against oxidative stress and mitochondrial DNA damage^76,79^. Similarly, the telomeric component TIN2, which together with TPP1 stabilizes the shelterin complex^80^ and recruits TERT to telomeres^81^, has been shown to shuttle from the nucleus to mitochondria in the context of enhanced oxidation to modulate mitochondrial respiration in favour of glycolysis^82^. Based on this knowledge, it would be interesting for future studies to follow the subcellular distribution of TERT and TIN2 (or other telomeric factors) as progenitors transition from naïve to primed pluripotency and determine their impact on telomere status and metabolic remodelling during this transition. Equally, it would be of interest to follow the behaviour of telomere trimming regulators, such as TZAP (telomeric zinc finger-associated protein), since in addition to being a negative regulator of telomere length, TZAP has been recognized as a transcriptional activator regulating mitochondrial metabolism^83,84^.

In conclusion, our findings unveil tight co-ordination between metabolic remodelling and ZSCAN4-associated telomere safeguarding in pluripotent progenitors ahead of differentiation and all subsequent development. As further evidence for the fundamental significance of telomere protective mechanisms, ESCs and adult stem cell progenitors with critically short telomeres have been shown to have impaired developmental potentials^63,72,85^. Given that some cancer cells express high levels of ZSCAN4 and the importance of metabolism reprogramming in tumour development, it is possible that metabolism-induced telomere regulation analogous to the one we discovered in this study might be hijacked in pathological conditions. Studying metabolic switches and telomere biology as two closely interrelated processes will thus be likely relevant for both fundamental and clinical research.

## Acknowledgments

We thank Taiping Chen and Wolf Reik for providing the pCAG-3XFlag-Zscan4c and pCAG-Zscan4c::eGFP plasmids, respectively. Thanks to Sandra Riad and King Hang Tommy Mau for help with the initial characterization of LD clones, to Danielle Admiraal, Dick W. Zijlmans and Pascal W.T.C. Jansen for help with ChEP, to Ruth Knops for help with 5mC quantification by LC-MS, to Laura Wingens for help with single-cell RNA-sequencing at the Sequencing Facility at the Radboud Institute for Molecular Life Sciences, to Chad Whilding and Dirk Dormann at the MRC LMS Microscopy Facility, and to James Elliot at the LMS/NIHR Imperial Biomedical Research Centre Flow Cytometry Facility. Gratitude also goes to Alice Jouneau, David Carling, Teresa Rayon, and Tristan Rodriguez for discussions and/or critical reading of the manuscript, and to all members of the Epigenetics and Development group. This work was supported by Genesis Research Trust – UK to V.A.; Biotechnology and Biological Sciences Research Council – UK to V.A. (BB/P005179/1) and to M.C. (BB/P005209/2; BB/H020233/1); the Dutch research organization – the Netherlands to H.M. (NWO XS - OCENW.XS22.3.119 and NWO XL - OCENW.XL21.XL21.100); Medical Research Council and UKRI Future Leaders Fellowship – UK to M.P. (MRC, MC_UP_1605/4 and MC_EX_MR/S015930/1); Imperial College President PhD scholarships – U.K. (D.K. and R.A.d.S.); Tommy’s National Miscarriage Centre – UK (C.L.N.); Babraham Institute – UK (B.d.C.S. and A.F.L-C.); Nottingham Trent University – UK (M.C.) and Imperial College London – UK (V.A., M.C, H.C.K., and M.B.).

## Methods

### Cell culture

Mouse ESCs were obtained from The Global Bioresource Center, ATCC (www.atcc.org), maintained at 37°C, 5% CO_2_ and routinely tested negative for mycoplasma contamination. ESCs were cultured on 0.1% gelatin-coated culture plates in GMEM-BHK12 basal medium (Gibco, 21710-025) supplemented with 10% foetal bovine serum (FBS) (Gibco, 10270-106), Leukaemia Inhibitory Factor (LIF) (made in-house), 0.1 mM β-mercaptoethanol (Gibco, 31350010), 0.25% (w/v) sodium bicarbonate (Gibco, 25080094), 1 mM sodium pyruvate (Gibco, 11360070), 0.1 mM non-essential amino acids (Gibco, 11140050), 2 mM L-glutamine (Gibco, 25030149), and 25 U/ml penicillin/streptomycin (Gibco, 11548876). For 2i/LIF conditions, ESCs were cultured on 0.2% gelatin-coated culture plates in DMEM F12: Neurobasal (vol. ratio 1:1) medium (Gibco, 11320033 and 21103049) supplemented with N2 (Gibco, 17502048), B27 (Gibco, 17504044), 0.1 mM β-mercaptoethanol, 2 mM L-glutamine, and 25 U/ml penicillin/streptomycin, with inhibitors CHIR99021 (3 µM) (Tocris, 4423), PD0325901 (1 µM) (Tocris, 4192) and LIF. For Knockout^TM^ serum replacement (KSR) (Gibco, 10828-028) or AlbuMAX (ThermoFisher, 11020021) treatment, cells were seeded the day before, followed by replenishing medium with FBS replaced by either 15% KSR or 10 % FBS supplemented with 1% AlbuMAX.

### Isolation of lipid droplet ESCs clones

E14 ESCs were plated at low density (1 x 10^4^ cells in a 10 cm dish) in FBS/LIF medium. After 8 days, individual colonies were isolated based on the presence or the absence of large LDs using phase-contrast microscopy. All clones were then expanded in FBS/LIF medium and frozen at early passages.

### Proliferation and self-renewal assays

To assess proliferation of ESC cultures, cells were seeded at 5 x 10^5^ cells per well. After 48 hours, cells were dissociated, counted and 5 x 10^5^ cells were re-plated. This was repeated for 7 passages and the growth curve was determined. To assess self-renewal capacity, alkaline phosphatase (AP) activity was measured using an AP staining kit (Sigma-Aldrich, 86R-1KT). ESCs were plated at 1 x 10^3^/ml, with medium replenished daily. Cells were grown for 7 days and stained according to the manufacturer’s instructions. Stained colonies were counted and scored as undifferentiated, mixed, and differentiated. At least 30 colonies were counted per sample.

### 3D spheroid differentiation

*In vitro* differentiation of 3D spheroids was carried out as previously described for imaging and molecular analyses^5,15^. Prior to setting up the experiment, ESCs were grown in routine culture. At 60% confluency, cells were then counted, and the required cell volume was washed in ice-cold phosphate-buffered saline (PBS). For imaging experiments, cells were resuspended in ice-cold Matrigel (Corning, 354230) by pipetting to a single cell suspension, at a density of 1500 cells/μl. 25 μl of this cell suspension was plated into an 8-well chamber slide and allowed to set at 37°C for 10 minutes. Pre-warmed differentiation medium (routine GMEM-BHK21 supplemented media with 15% FBS, and no LIF) was then added to the wells. Cells were subsequently fixed 24 hours or 48 hours after plating. For molecular analyses, plates were coated with ice-cold Matrigel to completely cover the well and allowed to set for 20 minutes at 37°C. For these experiments, cells were plated at a density of 5 × 10^4^ cells/cm^2^ in differentiation medium, supplemented with 5% Matrigel. Cells were collected at 24 hours or 48 hours after plating using Cell Recovery Solution (Corning, 354253) and cold PBS.

### Transfection

Cells were plated in a 6-well plate at a density of 2 x 10^5^ cells/well in routine medium. The next day, transfection medium was prepared consisting of 100 μl OptiMEM, 1 μg of plasmid DNA, and 1 μl of Lipofectamine 2000 (Thermo Fisher, 100014469). This complex was left at room temperature for 10 minutes. The plate was replenished with fresh 1.5 ml routine medium, followed by the addition of 100 μl of transfection medium and left overnight. The next day, medium was replaced with medium containing the appropriate antibiotic for selection. For stable cell lines, selection was carried out for 10 days.

### Mouse embryos *in vitro* culture

Mouse experiments were performed under a UK Home Office Project Licence. 4–6-week-old female and 2–6-month-old male CD1 mice (Charles River UK) were housed under 12-hour light:dark cycle and provided with ad libitum food and water in individually ventilated cages. Female mice were super-ovulated by intraperitoneal injection of 5 IU pregnant mare serum gonadotropin (PMSG; Folligon, MSD Animal Health), followed 46-48 hours later by 5 IU human chorionic gonadotropin (hCG; Chorulon, MSD Animal Health). Females were placed with males immediately following hCG injection and checked for copulatory plugs the following morning. Embryonic day (E) 2.5 embryos were collected from plugged females ~ 72-74 hours post hCG by flushing of dissected uteri with M2 medium (Sigma-Aldrich, M7167). Collected embryos were washed through successive drops of M2 and KSOMaa (Sigma-Aldrich, MR-106-D) pre-equilibrated at 37°C, 5% CO_2_. Following collection, embryos were cultured for 1 hour in KSOMaa at 37°C, 5% CO_2_ before either fixation (see Immunofluorescence and LD staining) or transfer to KSOMaa containing either DMSO (2% v/v as vehicle control;) or DGAT1/2i (25 or 50 µM each of two inhibitors; T863 for DGAT1 (Sigma-Aldrich, SML0539) and PF06424439 for DGAT2 (Tocris, 6348) and cultured for a further 24 hours, before fixation and processing for immunofluorescence and LD staining.

### Immunofluorescence and LD staining

Samples were washed and fixed in 4% paraformaldehyde (PFA; Sigma-Aldrich, P6148) for 10 minutes (self-renewal) or 20 minutes (ESC spheroids) at room temperature. Fixed samples were washed twice with PBS, followed by combined blocking and permeabilising with 0.3% TritonX-100 (Sigma-Aldrich, X100) in blocking buffer (3% bovine serum albumin (BSA; VWR, 422371X) in PBS) for 25 minutes at room temperature. The samples were then washed with PBS twice and incubated at 4°C overnight with the primary antibody in blocking buffer. The day after samples were washed twice and incubated with the secondary antibody, in blocking buffer, for 1 hour at room temperature. For staining of F-ACTIN, phalloidin (0.5 mg/ml; Sigma-Aldrich, P1951) was added to the secondary antibody mix and the incubation was carried out overnight at 4°C. All antibodies and dilutions are listed in Supplemental Methods Table 2.

For LD staining, samples were washed and incubated with BODIPY 493/503 (5 μg/ml in PBS; Invitrogen, D3922) for 25 minutes at room temperature, after the secondary antibody incubation. For nuclei staining, samples were washed and then incubated with DAPI (1 μg/ml in PBS; Sigma-Aldrich, D9542) for 10 minutes at room temperature. Samples were then mounted in ProLong Gold mounting agent (self-renewal) on slides or in 90% glycerol (ESC spheroids) for confocal microscopy. Confocal images were acquired at 1 µm z-sections, using a Leica SP5 microscope with a 63x oil immersion objective.

DAPI linescan analysis was performed as previously described^52,53^. ESCs were plated on coverslips in routine FBS/LIF culture conditions and fixed at 50% confluency. Coverslips were stained for DAPI and mounted on slides. Images were collected on a Leica SP5 confocal microscope with a 63x oil immersion objective. Low laser power of the 405 nm laser was used to prevent saturation and ensure distinct heterochromatin foci could be identified.

Embryos were fixed in 4% PFA in PBS for 20 minutes, washed 3 times for 10 minutes in PBS-PVA (0.1% PVA (Sigma-Aldrich, P8136-250G) in PBS), permeabilised in saponin-BSA for 1 hour, and blocked in 0.3% BSA-PBS for 1 hour. Samples were incubated in primary antibodies (1:100 anti-Zscan4, see Supplemental Methods Table 2) in saponin-BSA overnight at 4°C. The following day, samples were washed 3 times for 10 minutes in PBS-PVA, and incubated with secondary antibody (1:200, Alexa-Fluor555-anti rabbit) in saponin-BSA for 2 hours. Samples were washed once with PBS-PVA and incubated for 30 minutes in PBS-PVA containing BODIPY493/503 (5 mg/ml), DAPI (5 µg/ml) and Phalloidin-iFluor 647 (1:250; Abcam, ab176759). Samples were washed 3 times for 10 minutes in PBS-PVA, and mounted in Vectashield (2BScientific, H-1000). Confocal images were acquired at 1 µm z-sections encompassing the entire embryo, using a Leica Stellaris microscope with a 40x oil immersion objective.

### RNA extraction and RT-qPCR

RNA extraction was carried out using the RNeasy Mini kit (Qiagen, 74106) as per the manufacturer’s instructions. For reverse transcription (RT), 0.5-1 μg of RNA was mixed with 1 μl of oligo-dT and 1 μl of dNTPs, and then diluted with nuclease-free water up to 11 μl. Samples were incubated at 65°C for 5 minutes. Samples were then cooled on ice for 1 minute. To each sample, 1 μl of the enzyme SuperScriptIII reverse transcriptase (Invitrogen, 18080044), 1 μl of RNase inhibitor RNaseOut (Invitrogen, 10777019), 1 μl of supplied DL-dithiothreitol (DTT) and 4 μl of the supplied reaction buffer was added. The mixture was incubated at 50°C for 60 minutes, followed by incubation at 70°C for 15 minutes. The resultant cDNA was then diluted to a concentration of 5 ng/μl. This cDNA was then used for quantitative real-time polymerase chain reaction (PCR) or RT-qPCR using the StepOnePlus™ Real-Time PCR System (Applied Biosystems). All reactions were run in duplicate or triplicate in the same plate. The PCR program was as follows: 95°C for 15 minutes, followed by 45 cycles at 94°C for 15 seconds, 60°C for 30 seconds and 72°C for 30 seconds. For analysis, Ct values of analysed genes were normalized to the average of C_t_ values of the two housekeeping genes *L19* and *S17* (see Supplemental Methods Table 3 for primers).

### RNA-sequencing

Library preparation and sequencing was carried out on extracted RNA samples by Source Bioscience (UK) using Illumina HiSeq (2 x 150 bp) to a depth of 30 million reads. Raw FASTQ data was provided, and an initial quality check was performed using FastQC (Simon Andrews, Babraham Institute, Cambridge). The raw data was trimmed using Trimmomatic-0.33^86^ to remove Illumina adaptors and low quality bases (< 15). Resulting trimmed sequences were mapped to the mouse reference transcriptome mm10 (obtained from Ensembl) and quantified simultaneously by Kallisto (v0.43.1)^87^. Subsequent analysis was carried out in R/Bioconductor. Differential expression analysis was carried out using DESeq2 (v1.22.2)^88^, with lowly expressed transcripts filtered out (counts per million < 2 across more than 3 samples). Differentially expressed genes with a log2 fold-change > 1 and an adjusted p-value < 0.05 (following FDR correction) were used for subsequent over-representation analysis (ORA). WebGestalt 2019^89^ was used for the analysis in this work, using Benjamini-Hochberg (BH) as a method to control for false discovery rate (FDR) with FDR < 0.05 as a cut-off. Three databases were used: Reactome^90^, WikiPathways^91^, and Panther^92^. The raw RNA-seq data reported in this study have been deposited in Gene Expression Omnibus (GEO) under GSE271241.

### Single-cell RNA-sequencing

Cells were cultured in FBS/LIF conditions as mentioned previously. All cultures were passaged simultaneously at similar seeding density and grown for 72 hours before being processed on a 10X Genomics platform. For harvesting, cells were washed with DPBS and dissociated with 0.05% Trypsin-EDTA (Thermo Fisher, 25300054) for 5 minutes at 37°C in a humidified incubator. Dissociated cells were collected in 8 ml pre-warmed culture medium and centrifuged at 300 x g for 5 minutes. Cells were resuspended in pre-warmed culture medium and filtered through a 70 µm cell strainer. Filtered cells were centrifuged at 300 x g for 5 minutes and washed once with DPBS-BSA (DPBS with 0.04% BSA). Washed cells were resuspended in DPBS-BSA and filtered through a 40 µm cell strainer to obtain single cell suspensions. During further processing, cells were kept on ice. The cell concentrations of the single cell suspensions were determined using a haemocytometer and adjusted to 1 x 10^6^ cells/ml using DPBS-BSA. Single cell RNA-seq libraries were prepared from the single cell suspensions using a 10X Genomics Chromium Next GEM Single Cell 3’ Reagent Kit and 10X Chromium single-cell platform according to the manufacturer’s instructions (Doc. No. CG000204, revision B to C). DNA concentrations were assessed after cDNA amplification using a Qubit 1x dsDNA HS assay kit (Invitrogen, Q33231) and spectrophotometer (DeNovix ds-11). The DNA concentrations of the final libraries were determined in similar fashion. The peak size distributions of the final libraries were measured before pooling using a 2100 Bioanalyzer (Agilent) in combination with high sensitivity DNA Chips and Reagents (Agilent, 5067- 4626). The final library pools were sequenced on a NextSeq 500 (Illumina) sequencing instrument using a NextSeq 500/550 High Output 75-cycli kit (Illumina) with the following run type: 26bp read-1, 56bp read-2 and an 8bp index.

The sequencing data were processed and mapped against the 10X Genomics pre-built mouse reference transcriptome (mm10-2020-A) using 10X Genomics Cell Ranger (v6.1.2). The resulting raw count matrices were further analysed in R. Briefly, the DropletUtils package (v1.16.0) was used to perform barcode ranking (DropletUtils::barcodeRanks with ‘lower = 2500’ and ‘fit.bounds = 8000 - ∞’) and filter out barcodes of empty droplets (DropletUtils::emptyDrops with ‘lower = 7000’). The retained barcodes were filtered using Seurat (v4.3.0.1) (with 2100-3000 < detected genes < 9000 and raw transcript counts > 7500). Barcodes with more than 15-15.5% mitochondrial counts were removed. Additional filtering was performed with DropletQC (v0.0.0.9000) that identifies (and removes) damaged cells based on total RNA content and fraction of unspliced nuclear RNA (DropletQC::identify_damaged_cells with ‘nf_sep = 0.10’ and ‘umi_sep_perc = 50’). The count matrices were normalized individually using Seurat::SCTransform while regressing for the percentage of nuclear RNA and percentage of ribosomal protein counts. The count matrices were then concatenated with Seurat::merge. Initial downstream analyses included graph-based cell clustering using Seurat::FindClusters and Seurat::RunUMAP after determining a suitable clustering resolution using Clustree (v0.4.4). Normalized (and scaled) expression values computed with Seurat::SCTransform were used to generate violin and dot plots, unless stated otherwise. Gene signatures were calculated as module scores from Seurat::SCTransform-processed expression data using UCell (v2.0.1). The gene panels for calculating signature scores were: Naïve score; *Dazl, Esrrb, Klf2, Klf4, Nr0b1, Prdm14, Tbx3, Tcl1, Tfcp2l1, Zfp42* – Primed score; *Dnmt3a, Dnmt3b, Fgf5, Gata6, Krt18, Lef1, Lefty1, Lefty2, Oct6, Otx2, Sox17, Wnt8a, Zfp281* – Lipid score; *Acaca, Acly, Bscl2, Cidea, Dgat1, Dgat2, Fabp3, Fasn, G0s2, Gpat4, Ldlr, Scd1, Scd2, Srebf1, Srebf2, Tmem159*.

### Dimensionality reduction and clustering analysis

Accession numbers of published datasets and newly generated datasets used in this analysis are listed in Supplemental Methods Table 1. Expression matrices containing raw count data were generated from FASTQ files, using Kallisto^87^ (as described above). To minimise batch effects across the different studies, the count data from each of the datasets was pre-processed using ComBat^93^. These were then log transformed and a pre-filtering step was carried out to identify genes with a large variance across the samples, by setting thresholds for the minimum and maximum expression values of these genes. This resulted in a set of 6973 genes, which was further reduced to 2000 genes by iteratively optimising the projection and feature (i.e., gene) selection using a Gaussian process regression model^36^ in MATLAB. This low dimensionality clustering algorithm was used to generate the differentiation trajectory.

### Western blot

Samples were collected and lysed in RIPA buffer, supplemented with protease and phosphatase inhibitors, for protein extraction. Protein concentration was determined by BCA assay (ThermoFisher, 23225) as per the manufacturer’s instruction. 10-20 μg protein was loaded with Lammeli buffer (Bio-Rad, 161-0747) into SDS-PAGE gels (8-13% running gel/4% stacking gel) with pre-stained protein ladder. After SDS-PAGE, proteins were transferred to a methanol-activated PVDF Immobilon-FL membrane (Millipore, IPL00010) by semi-dry or wet transfer protocol. Blocking was carried out in 5% milk in Tris-buffered saline with 0.1% Tween-20 (TBST) for 1 hour at room temperature and then incubated in primary antibody solution (in 5% milk or BSA) overnight at 4°C. The membrane was washed in TBST, followed by incubation with a Horseradish peroxidase (HRP)-conjugated secondary antibody for 1 hour at room temperature. The membrane was washed in TBST, developed with Immobilon Forte HRP substrate (Bio-Rad, 1705061) and imaged by ImageQuant LAS4000 imager or exposed to X-ray film. Quantification was done in Fiji ImageJ. List of all antibodies and dilution factors used can be found in Supplemental Methods Table 2.

### Flow cytometry

Cells were grown in the specified culture conditions, dissociated with trypsin, and collected. For E14 ZE3-GFP reporter cells, cell pellets were directly resuspended in flow buffer (1 mM EDTA, 25 mM HEPES pH 7.0, 1% FBS). For ROS measurements, cells were stained with DCFDA (20 μM; Sigma-Aldrich, D8663) according to the manufacturer’s instructions in PBS. After washing, cells were resuspended in buffer. All experiments were carried out with an unstained control. For apoptosis measurements, eBioscience AnnexinV-FITC Apoptosis Detection Kit (Thermo Fisher Scientific, BMS500Fi) was used as per the manufacturer’s instructions. Flow cytometry experiments were carried out with BD FACSCalibur (Cell Quest software) or BD LSRII (FACSDiva software). Excitation lasers and emission filters were chosen as per manufacturer’s specifications and available lasers. Results were analysed in FlowJo (version 10.7.1).

### Seahorse assays

Seahorse assays for OCR and ECAR measurements were performed on an Agilent XFe24 (LD clones) or XFe96 analyser (CIDEA OE, ZSCAN4 OE). Cartridges were prepared as per the manufacturer’s instructions. Optimal cell density was determined for each plate (24-well or 96-well). Growth curve was calculated for each cell line prior to all experiments to ensure similar growth rates. For initial normalisation, cell number was calculated using DAPI fluorescence in the Seahorse plate. The cellular oxygen consumption rate (OCR) was measured via the Mitostress assay, with 3-5 cycles of measurement for each parameter. Basal respiration was measured first, prior to any treatment. To measure ATP-linked respiration, cells were treated acutely with oligomycin (1.5 μM; Sigma-Aldrich, O4876). This was followed by carbonyl cyanide 4- (trifluoromethoxy) phenylhydrazone (FCCP) (0.4 μM; Sigma-Aldrich, C2920) to measure maximum cellular respiration. Finally, antimycinA (2 μM; Sigma-Aldrich, A8674) and rotenone (2 μM, Sigma-Aldrich, R8875) were added in combination to determine non-mitochondrial respiration. The cellular extracellular acidification rate (ECAR) was measured via the Glycolysis stress assay, with cells starved of glucose prior to the assay. 3-5 cycles of measurement were performed for each parameter, starting with basal ECAR. Glucose (2.5 mM; Gibco, A2494001) was then added to the cells to determine glycolytic flux, followed by oligomycin (2.5 μM) to measure maximum glycolytic capacity. Finally, 2-deoxyglucose (2-DG) (50 mM; Sigma-Aldrich, D6134) was used to determine the non-glycolytic acidification rate. For substrate utilisation assays, cells were treated with specified inhibitors including UK5099 (20 μM; Tocris, 4186), BPTES (10 μM; Tocris, 5301), Etomoxir (40 μM, Sigma-Aldrich, E1905) for 1 hour prior to the assay, with continued treatment during the assay. This was carried out in conjunction with the Mitostress assay.

### Metabolomics Profiling

ESCs were plated at a density of 5 x 10^5^ cells/well in a 6-well plate, with two wells for each sample. After 16 hours in culture, the routine medium was changed to DMEM supplemented with 10% dialysed FBS for 24 hours. The media was aspirated, and the cells washed with cold (4°C) Ringer’s buffer, which was aspirated before addition of 750 μl of cold (−70°C) methanol. The methanol-quenched cells were then scraped from the well and the sample was transferred to a clean tube. To increase metabolite recovery, each well was washed with a further 750 μl of cold methanol and pooled with the first sample. The methanol-quenched samples were then dried in a rotary evaporator under reduced pressure. The dried cell pellets were frozen for subsequent metabolite extraction. Metabolites were extracted from the samples using a dual phase extraction method. To the frozen pellet, 300 μl of chloroform/methanol (2:1) was added and the samples vortexed for 30 seconds. An addition of 300 μl of HPLC-grade water was made and samples were vortexed for a further 30 seconds and centrifuged at 18,400 x g for 10 minutes. The upper aqueous and lower organic layers were transferred to separate silanized GC-MS vials. The extraction was repeated to maximise metabolite recovery. The organic fractions were dried under a stream of inert gas and stored at −80°C, while the aqueous fractions were freeze-dried and stored at −80°C. To each sample, 10 μl of myristic acid-d27 (1.5 mg/mL) was added as an internal standard, following which the samples were dried in a centrifugal evaporator. The dried aqueous samples were methoximated using 20 μl of methoxyamine (20 mg/mL) in anhydrous pyridine at 37°C for 90 minutes. This was followed by silylation with 80 μl of N-methyl-N-(trimethylsilyl) trifluoroacetamide (MSTFA) at 37°C for 30 minutes. Following derivatization, 2-fluorobiphenyl in anhydrous pyridine (10 μl, 1 mM) was added to the samples as an injection standard and the samples were vortexed, centrifuged for 5 minutes and transferred to clean silanized vials before GC-MS analysis. GC-MS analysis was performed as described below.

The dried apolar samples were reconstituted with a solution of anhydrous methanol/toluene (333μl, 1:1 v/v), treated with 0.5 M sodium methoxide (167 μl) and incubated at room temperature for 1 hour. The reaction was stopped by the addition of 1 M NaCl (500 μl) and concentrated HCI (25 μl). The apolar fraction was extracted with two volumes of hexane (500 μl) and the combined apolar layers were dried under inert gas. Samples were then silylated by reconstituting with 40 μl acetonitrile and treating with 40μl N-tert-Butyldimethylsilyl-N-methyltrifluoroacetamide (with 1% tert-Butyldimethyl-chlorosilane) (Sigma-Aldirch, 394882) and incubating at 70°C for 60 minutes. Following derivatisation, 2-fluorobiphenyl in anhydrous pyridine (10 μl, 1 mM) was added to the samples as an injection standard and the samples were vortexed, centrifuged for 5 minutes and transferred to clean silanized vials before GC-MS analysis. GC-MS analysis was performed as described below^94^.

### ^13^C_6_-glucose tracing experiment

ESCs were plated at a density of 2.5 x 10^5^ cells/well in a 6-well plate. After 16 hours in culture, the routine medium was changed to DMEM supplemented with 10% dialysed FBS and 5.6 mM unlabelled ^12^C_6_-glucose (Sigma-Aldrich, G8270) for 1 hour for equilibration. The media was then changed to DMEM with 10% dialysed FBS and 5.6 mM ^13^C_6_-glucose (Sigma-Aldrich, 389374), and the culture was maintained for 72 hours with no change of media. The cells were then washed, quenched and collected as described above for the unlabelled metabolomics study. Intracellular apolar metabolites (i.e. lipids) were extracted from six independent cultures of each clone analysed. To each sample, 10 μl of myristic acid-d27 (1.5 mg/mL) was added as an internal standard, following which the samples were dried in a centrifugal evaporator. Fatty Acid Methyl Esters (FAME) were generated by derivatisation with sodium methoxide (as above), which trans-esterifies all lipid bound fatty acids to a methyl ester. GC-MS was then performed as described below.

### Gas Chromatography-Mass Spectrometry (GC-MS)

The order of sample preparation and analysis on the GC-MS was randomized. The GC-MS analysis was performed using an Agilent Technology 7693 autosampler coupled to a 7980A GC system and a 5975 MSD mass spectrometer. 1ul of prepared sample was introduced into an inert glass splitless liner by the 7693 autosampler injector and chromatographically separated on a 30 m DB-5MS capillary column in the 7980A GC oven using helium as the carrier gas. The metabolites were detected via the 5975 MSD mass spectrometer running in electron impact ionisation mode. Metabolites were identified and assigned with in-house fragment/retention time databases or the Fiehn library in the deconvolution software AMDIS and individual peaks quantified using in-house MATLAB script based on the GAVIN3 software^94^. Mass isotopomers were quantified using in-house scripts (GAVIN) and the distributions were normalized so that the sum of the metabolite isotopomer abundances were equal to one and corrected for naturally occurring elemental isotopes based on previously described methods^95^.

### Liquid chromatography-mass spectrometry (LC-MS)

LC-MS was performed as previous described^96^. Briefly, lipids were extracted from frozen cells pellets using the Folch extraction method with chloroform/methanol/water (2:1:1). Lipids were dried using a SpeedVac (Thermo Fisher Scientific) and resuspended in chloroform/methanol (1:1). This was then injected into a Shimadzu Prominence 20-AD system (Shimadzu) for chromatographic separation with Waters Acquity UPLC C4 (300 Å, 1.7 μm particle size) columns. Using a mobile phase of water (solvent A) and acetonitrile (ACN) (solvent B) with 0.025% formic acid, 7 μl of the samples were eluted from the columns at 45°C. The flow rate was maintained at 100 μl/minute. Mass detection was carried out with an Orbitrap Elite mass spectrometer (Thermo Fisher Scientific), with an error cut-off of 5 parts per million (ppm). Source parameters for positive polarity: capillary temperature 275°C; source heater temperature 200°C; sheath gas 10 AU; aux gas 5 AU; sweep gas 5 AU. Source voltage was 3.8 kV. Full scan spectra in the range of m/z 340–1500 were acquired at a target resolution of 240,000 (FWHM at m/z 400). Xcalibur software (Thermo Fisher Scientific) was used for manual inspection of results, followed by data processing with Lipid Data Analyzer (LDA) 2.7.0_2019 software^97^.

### DNA methylation via mass spectrometry

DNA was isolated from cells using the PureLink Genomic DNA Mini Kit (Invitrogen, K182001) as per the manufacturer’s instructions. The samples were diluted to a final DNA concentration of 100 ng/μl, in a minimum volume of 3 μl. The DNA was degraded into individual nucleosides using DNA Degradase Plus (Zymo Research, E2020). Individual nucleosides were measured using a high-performance liquid chromatography-tandem mass spectrometry (HPLC-MS/MS) system. Quantification was performed using a standard curve derived from calibration standards. The 5mC and 5hmC levels were calculated as a concentration percentage ratio of % 5-methyl-2’-deoxycytidine to 2’-deoxyguanosine (%mdC/dG) and % 5-hydroxymethyl-2’-deoxycytidine to 2’-deoxyguanosine (%5hmdC/dG) respectively.

### Chromatin Enrichment for Proteomics (ChEP)

ChEP analysis was carried out as previously described^98^. Briefly, ESCs were grown in routine culture conditions, using T175 flasks. At least 3 x 10^6^ cells were used per replicate. Cells in flasks were washed with PBS and incubated in 1% formaldehyde (in PBS) at 37◦C for 10 minutes. Fixation was quenched using 0.25 M glycine at room temperature for 5 minutes. Cells were then washed with PBS twice and collected by scraping. Following centrifugation, cell pellets were frozen in liquid nitrogen. Cell pellets were lysed in lysis buffer, supplemented with protease inhibitors (Roche, 11697498001). The nuclei were pelleted via centrifugation and digested using lysis buffer containing 200 μg/ml RNase A (Thermo Fisher Scientific, EN0531). Centrifugation was repeated, and the nuclei were lysed in SDS buffer, with protease inhibitors, and urea buffer. The pellets were placed in storage buffer, supplemented with protease inhibitors, prior to sonication and centrifugation. The final supernatant, consisting of sonicated and cross-linked chromatin, was used to determine the protein yield via the Qubit Protein Assay kit (Invitrogen, Q33211). Cross-links were reversed with SDS-PAGE loading buffer. The protein mixtures were digested, desalted and LC-MS/MS was performed. MaxQuant software (v1.5.1.0) was used for analysis of MS data, with default settings to generate label-free quantification (LFQ) values for all samples. Perseus software (v1.5.0.15) was used for downstream analysis. Proteins were omitted if flagged as ‘potential contaminant’, ‘reverse hit’, or ‘only identified by site’. Proteins with less than two peptides were filtered out, and remaining LFQ values were log2 transformed. Proteins not detected in eight or more samples were excluded, and remaining missing values were imputed from a normal distribution using Perseus’ default settings (width=0.3 and down-shift=1.8). Fold changes and statistically different proteins were calculated for each comparison using a two-sample t-test.

### Telo-PCR

Genomic DNA was used to measure relative telomere length as previously described^99^. Briefly, genomic DNA was extracted from ESCs or spheroids using the DNeasy DNA extraction kit (Qiagen, 69504). qPCR was carried out using primers for either telomeric DNA or the single-copy acidic ribosomal phosphoprotein PO (36B4) gene (see Supplemental Methods Table 3). For each reaction 10 ng of DNA was used. Each PCR run also contained reactions, in triplicate, for a range of 3.75 to 60 ng of reference DNA (usually parental cell lines) for standard curve calculations. PCRs for telomeric DNA and 36B4 were run separately, as described below. For telomeric DNA PCR, each reaction contained 12.5 μl of SYBR Green qPCR ReadyMix, 300 nM of each primer, and nuclease-free water to make a 25 μl reaction, with the required DNA volume. The PCR program was as follows: 95°C for 10 minutes, followed by 30 cycles of 95°C for 15 seconds and 56°C for 1 minute. For 36B4 PCR, each reaction contained 12.5 μl of SYBR Green qPCR ReadyMix, 300 nM of the forward primer, 500 nM of the reverse primer, and nuclease-free water to make a 25 μl reaction with the required DNA volume. The PCR program was as follows: 95°C for 10 minutes, followed by 35 cycles of 95°C for 15 seconds, 52°C for 20 seconds and finally, 72°C for 30 seconds. To calculate relative telomere lengths, input amounts were determined using the standard curve. The ratio of relative input amount for telomeric DNA to 36B4 was then calculated and represented as average telomere length ratio across the samples.

### C-Circle amplification PCR

ALT-specific C-circles were measured as a readout of ALT activity as previously described^100^. Genomic DNA was extracted using the DNeasy DNA kit as per manufacturer’s instructions. C-circle amplifications (CCA) were carried out for each sample using a thermocycler. For each reaction, 30 ng of DNA was used with ф29 DNA Polymerase (7.5 units; New England Biolabs, M0269), dNTPs (1 mM each), Tween20 (0.1% v/v; Sigma Aldrich, P1379), DTT (1 mM), BSA (1 µg/ml; provided with enzyme), and ф29 buffer (provided with enzyme) in Tris buffer pH 7.6 (10 mM). A control reaction without ф29 DNA Polymerase was carried out for each sample. The amplification was performed as follows: 30°C for 4 hours, followed by 70°C for 2 minutes, and stored at 4°C. CCA was then followed by a qPCR reaction using the primers (telomeric DNA and single-copy 36B4) for telomere length analysis in Supplemental Methods Table 3. Each reaction contained 5 μl of the CCA product, with 12.5 μl of SYBR Green qPCR ReadyMix, 500 nM of each primer, and nuclease-free water to make a 25 μl reaction. For telomeric DNA PCR, a standard curve was calculated using serial three-fold dilutions to obtain the following concentrations (ng/μl) of parental cell line DNA: 3.2, 1.07, 0.36, 0.12, 0.04 and 0.013. The PCR program was as follows: 95°C for 15 minutes, followed by 30 cycles of 95°C for 7 seconds and 58°C for 10 seconds. For single-copy gene PCR a standard curve was calculated using serial three-fold dilutions to obtain the following concentrations (ng/μl) of parental cell line DNA: 3.2, 1.6, 0.8, 0.4, 0.2 and 0.1. The PCR program was as follows: 95°C for 5 minutes, followed by 40 cycles of 95°C for 15 seconds and 58°C for 30 seconds. ALT-positive controls were always run in parallel (here an osteosarcoma cell line, U-2 OS was used as recommended). For quantification, input DNA amounts were determined using the standard curve and normalised to single copy gene levels as described above. CCA levels were then determined by subtracting the amount of DNA of the respective control ф29-negative reactions and normalised to U-2 OS CCA levels. CCA-positive samples are those that show CCA level >1% of the CCA level for U-2 OS.

### Metaphase preparation and quantitative FISH

ESCs were treated with colcemid (0.05 μg/ml; Roche, 10295892001) for 90 minutes under routine culture conditions. After this, cells were washed with PBS, and dissociated, and pelleted. Cell pellets were lysed with 8 ml of pre-warmed hypotonic solution (8 g/l sodium citrate; Sigma-Aldrich C8532) added dropwise, and then incubated at 37°C for 20 minutes. To this, 1 ml of cold fixative (ethanol:acetic acid/3:1) was added and centrifuged at 200 x g for 5 minutes. The supernatant was discarded, more fixative was added to the cell pellet dropwise, and tubes were centrifuged at 300 x g for 5 minutes; fixation was repeated. Lastly, 8 ml of fixative was added and left overnight at −20°C. The following day, slides were immersed in 100% ethanol. Cells were thawed at room temperature for at least 30 minutes prior to centrifugation at 300 x g for 5 minutes. Cell pellets were resuspended in 1 ml of fixative. A humid environment was created using wet paper towels under the slides. On each slide, 50 μl of cell suspension was dropped to allow for good metaphase spreads. Slides were dried overnight at room temperature. The pre-dried metaphase spreads, 35 μl of hybridisation solution containing the TelC-probe (PNA Bio, USA, F1002) (1 nM probe in 70% w/v formamide, 20 mM Tris-Cl pH 7.4, 1% w/v BSA) was added per slide and denatured by incubation at 80°C for 3 minutes. This was followed by incubation at room temperature in the dark, for 2 hours. Following incubation, the slides were washed successively with the two wash buffers. First, with wash buffer 1 (70% w/v formamide, 20 mM Tris-Cl pH 7.4), twice for 15 minutes each time, then with wash buffer 2 (70% w/v formamide, 150 mM NaCl, 0.05% v/v Tween-20), thrice for 5 minutes each time. The slides were then dehydrated with successive ethanol baths (70%, 80%, 90%, 100%) and then air dried in the dark. Coverslips were then mounted with DAPI (5 μg/ml) and the slides were sealed. Confocal images were acquired using a Leica SP5 microscope with a 63x glycerol immersion objective.

### Image analysis

All image analysis was carried out using ImageJ/Fiji (v1.53p) and CellProfiler (v4.2.1)^101^. For BODIPY-based neutral lipid quantification in ESCs, custom macro scripts in ImageJ were used to quantify total area of BODIPY signal and total area of DAPI signal in a maximum z-stack projection. For quantification of ZSCAN4-positive (Z4+) ESCs, wide-field images were taken and converted to grayscale using custom macros. Then number of cells were quantified in each channel using Fiji’s ‘Analyse Particles’ plugin. For quantification of chromocenters using DAPI, lines were drawn across an individual nucleus to intersect with the maximum number of DAPI foci as shown in Figure 4G (insets). Intensity across the line was then measured using ImageJ/Fiji and the number of peaks was recorded. For telomere quantitative-FISH, CellProfiler pipelines were created to split channels and convert to grayscale. TelC-Cy3 intensity (telomere foci) was quantified within DAPI regions at chromosomal ends, for signal normalisation. For APB quantification, slices were extracted from 3D stacks and converted to grayscale using custom macros. These were then imported into CellProfiler to quantify foci signal within DAPI regions in each slice. Colocalised PML/TRF1 foci were then counted within the nuclear regions in each slice and normalised to total number of PML foci.

For embryo analysis, images were processed to single channel maximum projection or single-plane images using Fiji. Using DAPI and phalloidin staining, embryo morphology was manually scored as normal (morula or E3.5 blastocyst), abnormal, which included embryos with collapsed or multiple cavities, and embryos with apoptotic cells (see Figure S7 for examples and scoring). Apoptotic cells had extremely high BODIPY signal therefore these embryos were excluded from further analyses. The number of BODIPY foci per embryo were identified in maximum projection images using CellProfiler (v4.2.0). ZSCAN4 staining intensity was quantified, using CellProfiler, in single z-sections within segmented nuclei (see Figure S5D for overview of analysis pipeline). Several z-sections >=5 sections apart were analysed per embryo to avoid double-sampling the same nucleus in different focal planes.

### Statistical analysis

Statistical analyses were performed using GraphPad Prism (v10.0.2) except for bulk/single-cell RNA-sequencing and ChEP data. Data are presented as means ± standard error of mean (s.e.m.), unless otherwise specified in the figure legends. Number of replicates are stated in the figure legends. Statistical tests used for each set of experiments are specified in the figure legends; in this study data were compared using either a two-tailed paired or unpaired Welch’s t-test, or one- or two-way ANOVA with a post-hoc test as specified in text. For quantitative analyses, including quantification of Z4+ ESCs, APB formation, and embryo analysis, normality was tested using D’Agostino and Pearson’s normality test. For non-normally distributed data, the Kruskal-Wallis test with Dunn’s multiple correction test was used. No statistical method was used to predetermine sample size of embryos.

## Author contributions

R.A.d.S. carried out most experiments and data analyses; D.B., D.K. and M.P.B. generated and/or contributed to the characterization of LD clones under the expert guidance of M.C.; N.G.P.B. generated and analyzed the scRNA-seq data under the supervision of H.M.; Y.Z. generated and/or contributed to the characterization of ZSCAN4 OE and CIDEA OE ESCs; H.M. supervised the ChEP generation and analysis performed by E.J.E.M.K.; C.L.N. performed Telo-FISH on 3D spheroids and provided expert guidance on telomere analyses; H.C.K. supervised the GC-MS experiments performed by D.B., D.K., and J.K.E.; M.P. supervised the analysis of cultured pre-implantation embryos performed by B.J.L.; M.B. supervised the dimensionality reduction and clustering analysis performed by Z.L.; TAG quantifications by LC-MS were performed by B.C.S. and A.F.L-C; 5mC quantifications by LC-MS were performed by H.M. and J.H.J.; V.A. conceived the project and supervised the study; V.A. and R.A.d.S. wrote the manuscript with contributions from all the co-authors.

## Supplemental Information

Supplemental Figures: Figures S1-S6

Supplemental Methods Table 1: Excel file containing accession numbers for transcriptomic datasets, related to Figure 1K.

Supplemental Methods Table 2: Excel file containing antibodies and dilutions used.

Supplemental Methods Table 3: Excel file containing primers used for qPCR and telomeric analyses.

