## Supplemental Figures for "The exit of naïve pluripotency contains a lipid metabolism-induced checkpoint for genome integrity"

### FIGURE S1

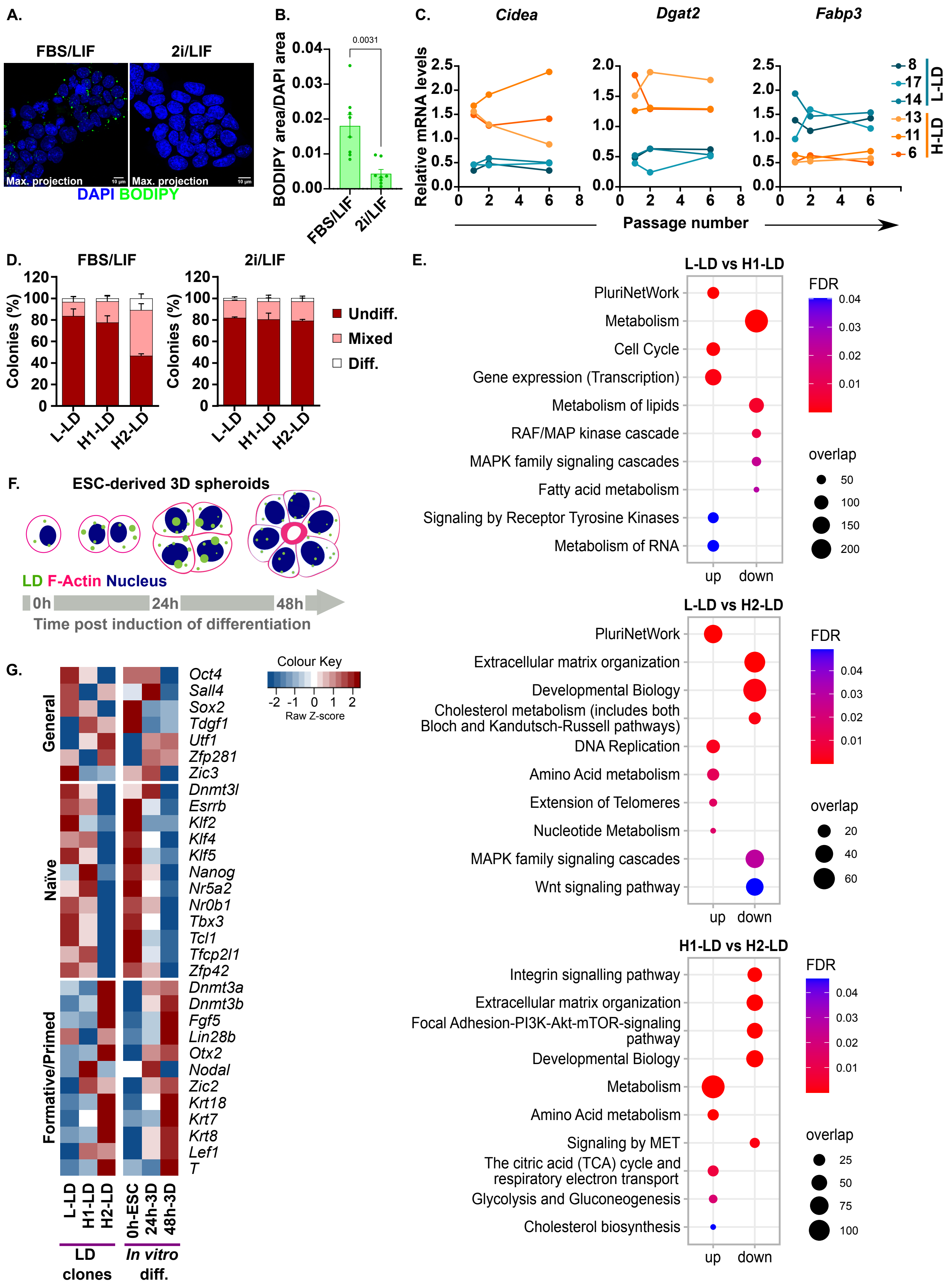

A.
FBS/LIF
2i/LIF
Max. projection
DAPI BODIPY
B.
BODIPY area/DAPI area
0.0031
FBS/LIF
2i/LIF
C.
Cidea
Dgat2
Fabp3
Relative mRNA levels
Passage number
L-LD
H-LD
D.
FBS/LIF
2i/LIF
Colonies (%)
Undiff.
Mixed
Diff.
L-LD
H1-LD
H2-LD
E.
L-LD vs H1-LD
PluriNetWork
Metabolism
Cell Cycle
Gene expression (Transcription)
Metabolism of lipids
RAF/MAP kinase cascade
MAPK family signaling cascades
Fatty acid metabolism
Signaling by Receptor Tyrosine Kinases
Metabolism of RNA
FDR
overlap
up
down
L-LD vs H2-LD
PluriNetWork
Extracellular matrix organization
Developmental Biology
Cholesterol metabolism (includes both Bloch and Kandutsch-Russell pathways)
DNA Replication
Amino Acid metabolism
Extension of Telomeres
Nucleotide Metabolism
MAPK family signaling cascades
Wnt signaling pathway
H1-LD vs H2-LD
Integrin signalling pathway
Extracellular matrix organization
Focal Adhesion-PI3K-Akt-mTOR-signaling pathway
Developmental Biology
Metabolism
Amino Acid metabolism
Signaling by MET
The citric acid (TCA) cycle and respiratory electron transport
Glycolysis and Gluconeogenesis
Cholesterol biosynthesis
F.
ESC-derived 3D spheroids
LD F-Actin Nucleus
0h
24h
48h
Time post induction of differentiation
G.
Colour Key
Raw Z-score
General
Naïve
Formative/Primed
Oct4
Sall4
Sox2
Tdgf1
Utf1
Zfp281
Zic3
Dnmt3l
Esrrb
Klf2
Klf4
Klf5
Nanog
Nr5a2
Nr0b1
Tbx3
Tcl1
Tfcp2l1
Zfp42
Dnmt3a
Dnmt3b
Fgf5
Lin28b
Otx2
Nodal
Zic2
Krt18
Krt7
Krt8
Lef1
T
0h-ESC
24h-3D
48h-3D
LD clones
In vitro diff.

**Figure S1: L-LD and H-LD phenotypes correspond to distinct pluripotent states in ESCs, related to Figure 1.**

(A) Representative images of BODIPY 493/503-stained LDs (green) and DAPI-stained nuclei (blue) in E14-ESCs cultured in FBS/LIF or 2i/LIF. Images shown are z-stack maximum projections, scale bars, 10 μm.

(B) Quantification of BODIPY signal, normalized to DAPI (area). For each culture, 4 colonies were imaged in independent experiments (n=2). Error bars, means ± s.e.m.; two-tailed t-test with Welch's correction.

(C) Gene expression profiling (RT-qPCR) for lipid metabolism genes in L-LD (8, 14, 17) and H-LD (6, 11, 13) clones at passages 1, 2 and 6 post isolation from bulk E14-ESC culture.

(D) Percentage of colonies, in L-LD (8), H1-LD (4) and H2-LD (11), formed from cells seeded at low density and grown for 7 days in FBS/LIF (left) and 2i/LIF (right). Colonies were counted and scored as undifferentiated, mixed, and differentiated based on alkaline phosphatase staining. Error bars, means ± s.e.m. (n=3).

(E) Dot plots showing selected over-represented pathways identified using differentially expressed genes between L-LD and H1-LD (top panel), L-LD and H2-LD (middle panel), and H1-LD and H2-LD (bottom panel). Each dot represents a specific pathway, where the size of the dot corresponds to the number of differentially expressed genes associated with that pathway, and the colour of the dot corresponds to the false discovery rate (FDR) value following Benjamini-Hochberg (BH) correction for multiple testing (FDR < 0.05).

(F) Schematic representation of ESC-derived 3D spheroids and lipid storage trafficking prior to (0 hour), and at different timepoints post induction of differentiation (24 and 48 hours).

(G) Heatmap showing relative expression of pluripotency-associated genes in LD clones (left) and ESC-derived 3D-spheroids upon differentiation (right).

### FIGURE S2

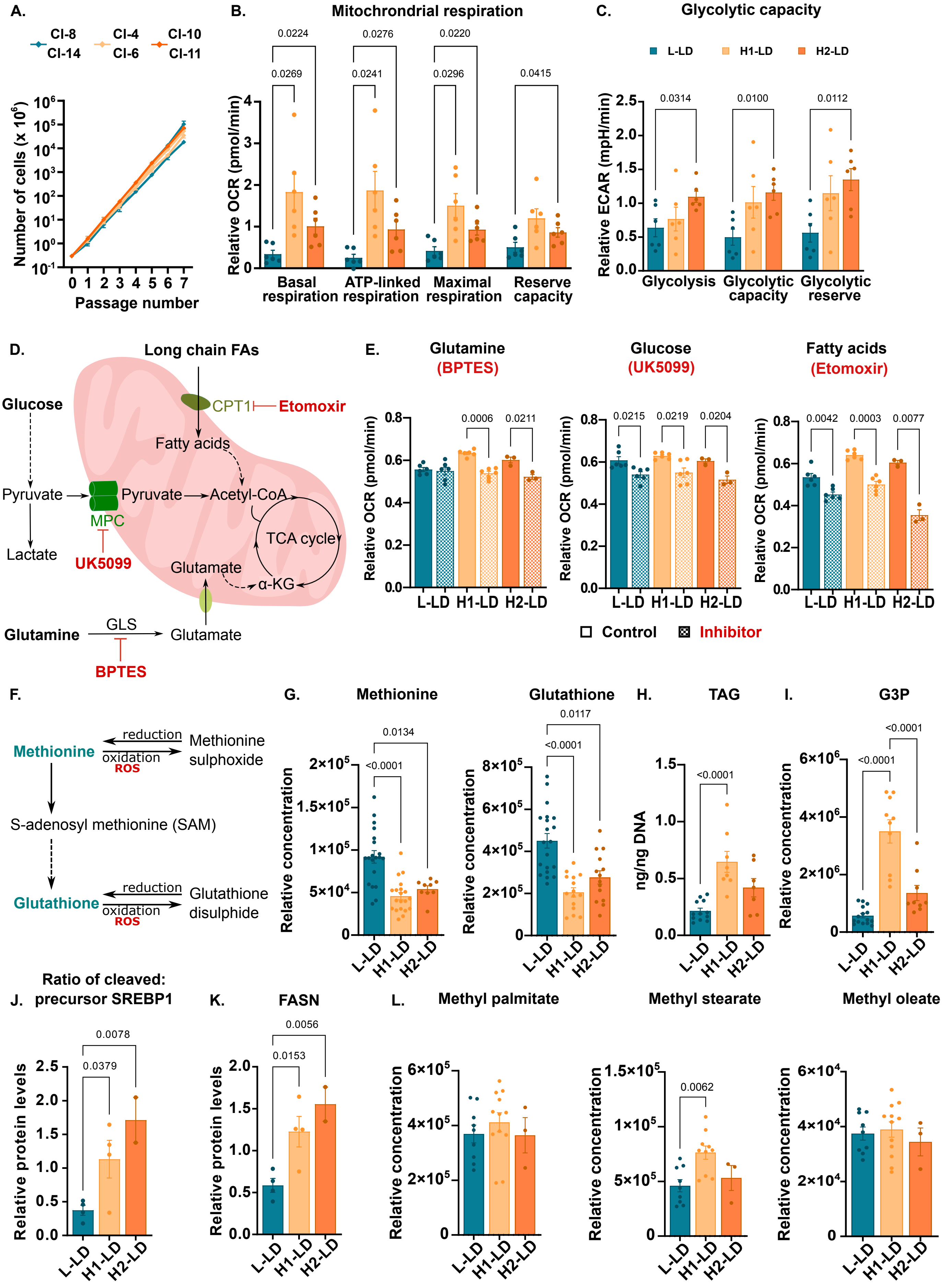
A.
Cl-8
Cl-14
Cl-4
Cl-6
Cl-10
Cl-11
Number of cells (x 10^6)
Passage number
B.
Mitochrondrial respiration
Relative OCR (pmol/min)
Basal respiration
ATP-linked respiration
Maximal respiration
Reserve capacity
C.
Glycolytic capacity
L-LD
H1-LD
H2-LD
Relative ECAR (mpH/min)
Glycolysis
Glycolytic capacity
Glycolytic reserve
D.
Long chain FAs
Glucose
CPT1
Etomoxir
Fatty acids
Pyruvate
Pyruvate
Acetyl-CoA
MPC
TCA cycle
Lactate
UK5099
Glutamate
α-KG
Glutamine
GLS
Glutamate
BPTES
E.
Glutamine (BPTES)
Glucose (UK5099)
Fatty acids (Etomoxir)
Relative OCR (pmol/min)
Control
Inhibitor
F.
Methionine
reduction
oxidation
ROS
Methionine sulphoxide
S-adenosyl methionine (SAM)
Glutathione
Glutathione disulphide
G.
Methionine
Glutathione
Relative concentration
H.
TAG
ng/ng DNA
I.
G3P
Relative concentration
J.
Ratio of cleaved: precursor SREBP1
Relative protein levels
K.
FASN
Relative protein levels
L.
Methyl palmitate
Methyl stearate
Methyl oleate
Relative concentration

**Figure S2: Metabolic characteristics of LD clones along a differentiation trajectory, related to Figure 2 and Figure 3.**

(A) Growth curves of specified LD clones in FBS/LIF conditions over 7 passages (14 days) (n=3).

(B) Assessment of mitochondrial respiration from OCR measurements following the serial addition of ETC-complex inhibitors in LD clones (see also Figure 2A).

(C) Assessment of glycolytic capacity from ECAR measurements following addition of indicated drugs (see also Figure 2B). Error bars, means ± s.e.m.; ANOVA with Tukey's post-hoc (n=3 per clone).

(D) Schematic representation of mitochondrial metabolic pathways with indication of inhibitors (red) used in (E).

(E) ATP-linked respiration to assess substrate utilization in L-LD (8, 17), H1-LD (4, 6) and H2-LD (11) clones treated with a glutaminase inhibitor (BPTES, 10 μM), MPC1 inhibitor (UK5099, 20 μM), and CPT1 inhibitor (Etomoxir, 40 μM), respectively. Error bars, means ± s.e.m.; paired two-tailed t-test (n=3).

(F) Schematic of methionine and glutathione redox metabolism.

(G) Relative concentration of methionine and glutathione in L-LD (5, 6, 14, 17), H1-LD (1, 4, 6, 13) and H2-LD (10, 11) measured by GC-MS. For each clone, 5 replicates were carried out; each data point represents one replicate of an individual clone. Error bars, means ± s.e.m.; ANOVA with Tukey's post-hoc (n=3 per clone).

(H) LC-MS quantification of TAG in L-LD (8,14,17), H1-LD (4, 6) and H2-LD (10, 11) (see also Figure 1D).

(I) Relative concentration of glycerol-3-phosphate (G3P) in L-LD (8, 14, 17), H1-LD (4, 6) and H2-LD (10, 11) measured by GC-MS (as shown in Figure 2D).

(J,K) Quantification of (J) cleaved to precursor SREBP1 and (K) FASN to $\alpha$-TUBULIN loading control in L-LD (8, 17), H1-LD (4, 6), and H2-LD (11) clones. Each dot represents one replicate. Error bars, means ± s.e.m.; ANOVA with Tukey's post-hoc test (n=2 per clone).

(L) Relative concentration of intracellular methylated fatty acids (methyl palmitate, methyl stearate, methyl oleate) in L-LD (8, 14, 17), H1-LD (1, 4, 6, 13), and H2-LD (11) clones. Each dot represents one replicate. Error bars, means ± s.e.m.; ANOVA with Tukey's post-hoc test (n=3 per clone).

### FIGURE S3

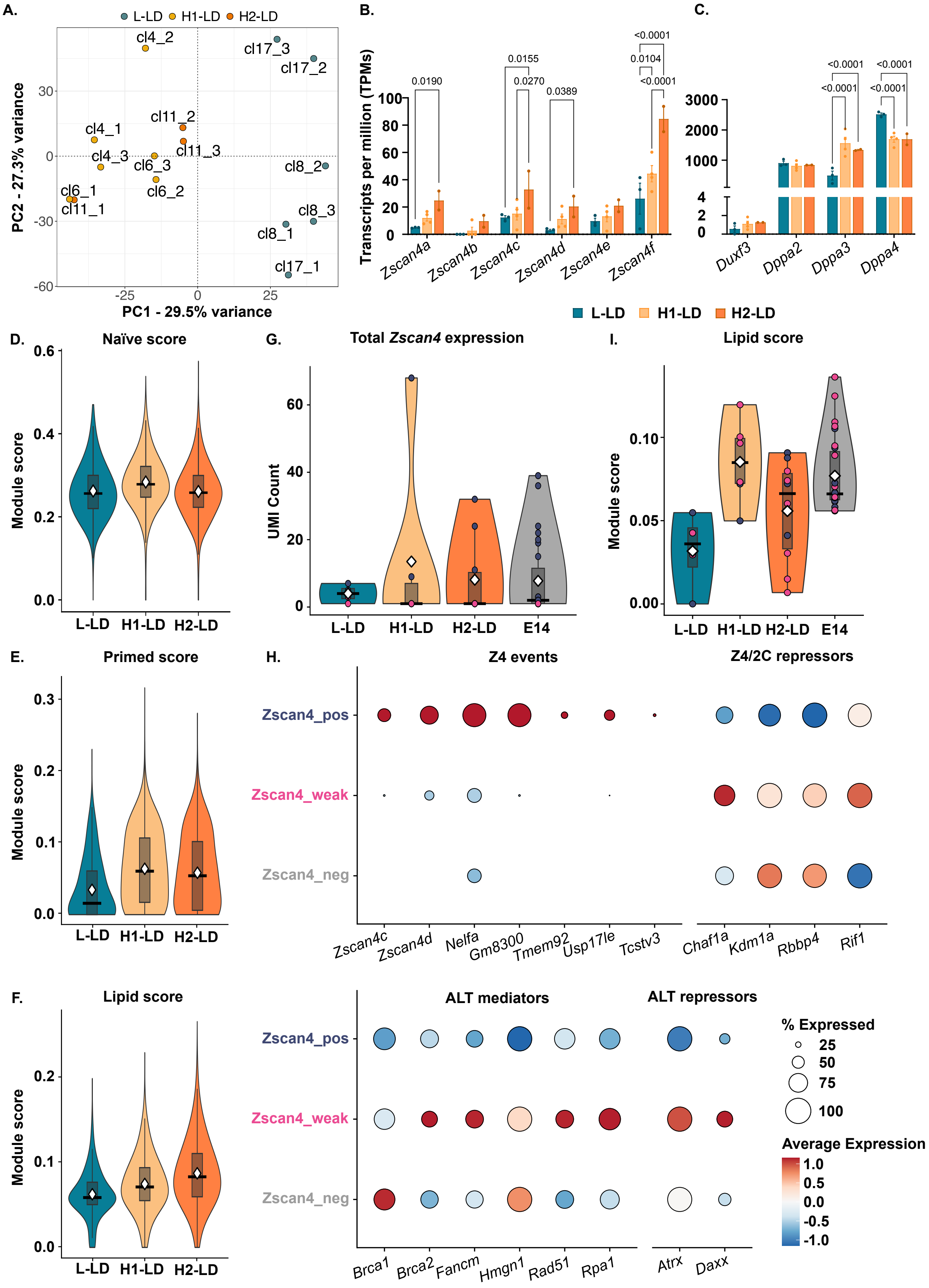

A.
L-LD H1-LD H2-LD
PC1 - 29.5% variance
PC2 - 27.3% variance
B.
Transcripts per million (TPMs)
Zscan4a
Zscan4b
Zscan4c
Zscan4d
Zscan4e
Zscan4f
C.
Duxf3
Dppa2
Dppa3
Dppa4
L-LD H1-LD H2-LD
D.
Naïve score
E.
Primed score
F.
Lipid score
Module score
G.
Total Zscan4 expression
UMI Count
L-LD H1-LD H2-LD E14
I.
Lipid score
H.
Z4 events
Z4/2C repressors
Zscan4_pos
Zscan4_weak
Zscan4_neg
Zscan4c Zscan4d Nelfa Gm8300 Tmem92 Usp17le Tcstv3
Chaf1a Kdm1a Rbbp4 Rif1
ALT mediators
ALT repressors
Brca1 Brca2 Fancm Hmgn1 Rad51 Rpa1
Atrx Daxx
% Expressed
Average Expression

**Figure S3: Multi-omics analysis reveals differential ZSCAN4 activity in LD clones, related to Figure 4.**

(A) PCA performed on ChEP data collected from L-LD (8, 17), H1-LD (4,6) and H2-LD (11) clones.

(B) Gene expression profiling (RNA-seq) of *Zscan4* paralogues in L-LD (8, 14, 17), H1-LD (1, 4, 6, 13) and H2-LD (10,11) clones represented as TPMs.

(C) Gene expression profiling (RNA-seq) of *Duxf3*, *Dppa2, Dppa3* and *Dppa4* in LD clones as above, represented as TPMs. Error bars, means ± s.e.m.; Fisher's two-way ANOVA.

(D,F) Pseudo-bulk analysis of single-cell RNA-seq data collected from representative L-LD (8), H1-LD (4) and H2-LD (11) clones to confirm distinct (D) naïve, (E) primed, and (F) lipid transcriptional signatures computed as scores (see Methods).

(G) Expression of all *Zscan4* paralogues in each clone group and parental E14-ESCs (E14) plotted as UMI counts.

(H) Expression of known genes associated with ZSCAN4 (Z4) events (*Zscan4c, Zscan4d, Nelfa, Gm8300, Tmem92, Usp17le, Tcstv3*), repressors of Z4 events (*Chaf1a, Kdm1a, Rbbp4, Rif1*), mediators of alternative lengthening telomere (ALT) mechanism (*Brca1, Brca2, Fancm, Hmgn1, Rad51, Rpa1*) and repressors of ALT (*Atrx, Daxx*) in *Zscan4*-negative, weakly expressing, and positive single cells shown as dot plots (see also Figure 4I).

(I) Lipid transcriptional score computed for *Zscan4*-expressing cells in each clone group and parental E14-ESCs.

### FIGURE S4

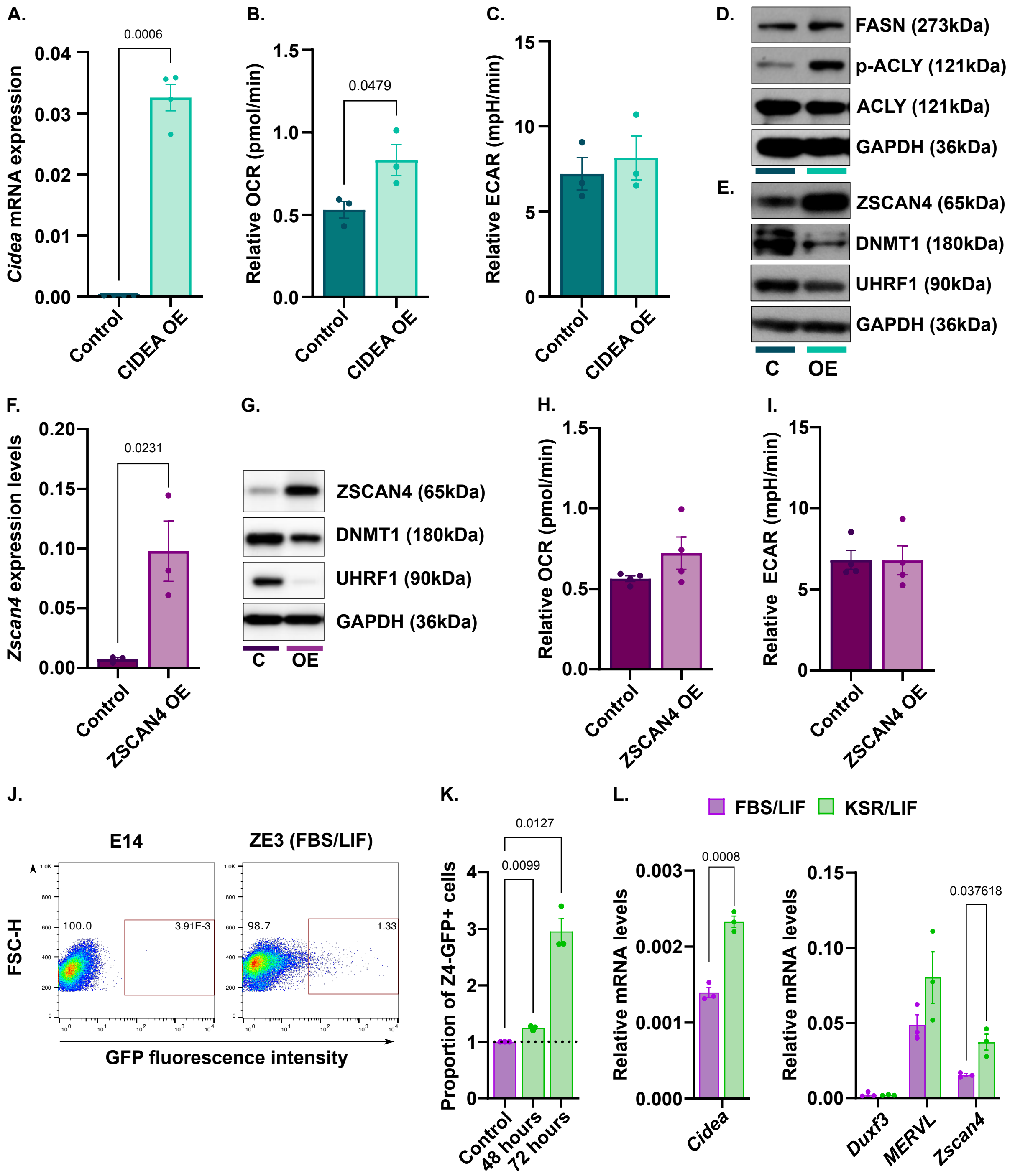
A.
0.0006
Cidea mRNA expression
0.04
0.03
0.02
0.01
0.00
Control
CIDEA OE
B.
Relative OCR (pmol/min)
1.5
1.0
0.5
0.0
0.0479
Control
CIDEA OE
C.
Relative ECAR (mpH/min)
15
10
5
0
Control
CIDEA OE
D.
FASN (273kDa)
p-ACLY (121kDa)
ACLY (121kDa)
GAPDH (36kDa)
E.
ZSCAN4 (65kDa)
DNMT1 (180kDa)
UHRF1 (90kDa)
GAPDH (36kDa)
C
OE
F.
Zscan4 expression levels
0.20
0.15
0.10
0.05
0.00
0.0231
Control
ZSCAN4 OE
G.
ZSCAN4 (65kDa)
DNMT1 (180kDa)
UHRF1 (90kDa)
GAPDH (36kDa)
C
OE
H.
Relative OCR (pmol/min)
1.5
1.0
0.5
0.0
Control
ZSCAN4 OE
I.
Relative ECAR (mpH/min)
15
10
5
0
Control
ZSCAN4 OE
J.
E14
ZE3 (FBS/LIF)
FSC-H
100.0
3.91E-3
98.7
1.33
GFP fluorescence intensity
K.
0.0127
0.0099
Proportion of Z4-GFP+ cells
4
3
2
1
0
Control
48 hours
72 hours
L.
FBS/LIF
KSR/LIF
0.0008
Relative mRNA levels
0.003
0.002
0.001
0.000
Cidea
Relative mRNA levels
0.15
0.10
0.05
0.00
0.037618
Duxf3
MERVL
Zscan4

**Figure S4: *Zscan4* induction is downstream of lipid storage in ESCs, related to Figure 5.**
(A) Expression levels (RT-qPCR) of *Cidea* transcript, normalised to housekeeping genes (*S17, L19*), in CIDEA OE and control ESCs. Error bars, means ± s.e.m.; paired two-tailed t-test with Welch's correction (n=4).
(B) ATP-linked OCR and (C) glycolysis-associated ECAR, measured by Seahorse analysis in CIDEA OE and control ESCs. Error bars, means ± s.e.m.; paired two-tailed t-test with Welch's correction (n=3).
(D, E) Representative western blots of (D) FASN, p-ACLY (Ser455) and total ACLY, with GAPDH loading control, and (E) ZSCAN4, DNMT1, UHRF1, with GAPDH loading control in CIDEA OE (labelled OE) and control (labelled C) ESCs (n=3).
(F) Expression level (RT-qPCR) of total *Zscan4*, normalised to housekeeping genes (*S17, L19*), in ZSCAN4 OE and control ESCs. Error bars, means ± s.e.m.; paired two-tailed t-test with Welch's correction (n=3).
(G) Representative western blots of ZSCAN4, DNMT1, UHRF1, with GAPDH loading control in ZSCAN4 OE and control ESCs (n=3).
(H) ATP-linked OCR and (I) glycolysis-associated ECAR, measured by Seahorse analysis in ZSCAN4 OE and control ESCs. Error bars, means ± s.e.m.; paired two-tailed t-test with Welch's correction (n=4).
(J) Representative scatter plots to depict flow cytometry gating strategy for Zscan4c::GFP (E14-ZE3) reporter ESC line, using parental E14-ESCs as a negative-GFP control. (K) Proportion of ZSCAN4::GFP positive (Z4-GFP+) ESCs in FBS/LIF (control) or KSR/LIF conditions for 48 hours and 72 hours.
(L) Relative expression of *Cidea, Duxf3, MERVL*, and total *Zscan4*, normalised to housekeeping genes (*S17, L19*) in ESCs cultured in FBS/LIF or KSR/LIF conditions for 48 hours. Error bars, means ± s.e.m.; paired two-tailed t-test with Welch's correction (n=3).

#### FIGURE S5

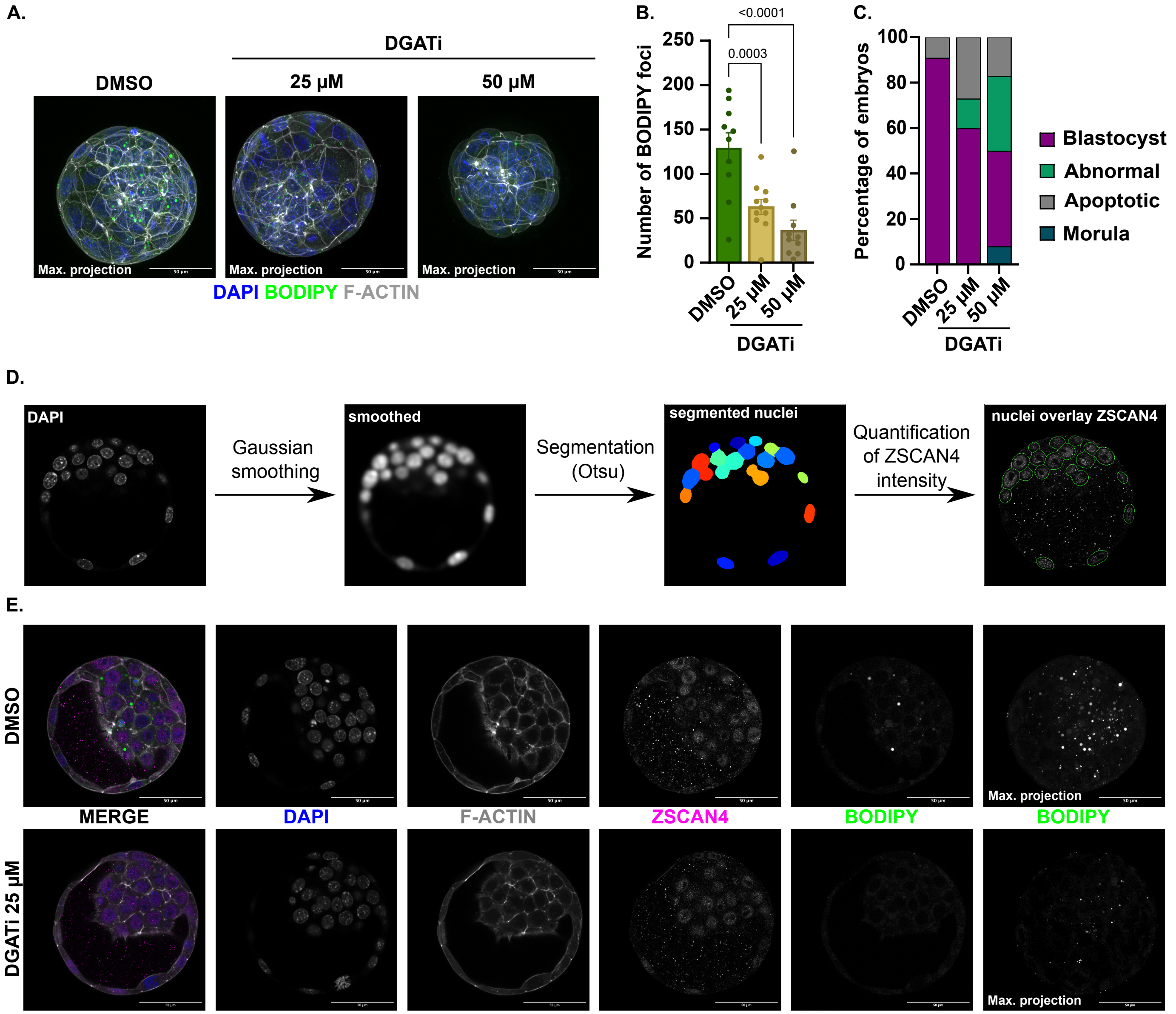
A.
DGATi
DMSO
25 µM
50 µM
Max. projection
DAPI BODIPY F-ACTIN
B.
<0.0001
0.0003
Number of BODIPY foci
250
200
150
100
50
0
DMSO
25 µM
50 µM
DGATi
C.
Percentage of embryos
100
80
60
40
20
0
DMSO
25 µM
50 µM
DGATi
Blastocyst
Abnormal
Apoptotic
Morula
D.
DAPI
Gaussian smoothing
smoothed
Segmentation (Otsu)
segmented nuclei
Quantification of ZSCAN4 intensity
nuclei overlay ZSCAN4
E.
DMSO
DGATi 25 µM
MERGE
DAPI
F-ACTIN
ZSCAN4
BODIPY
BODIPY
Max. projection

**Figure S5: Interfering with lipid storage during the morula-to-blastocyst transition reduces ZSCAN4 expression *in vivo*, related to Figure 5.**

(A) Representative maximum projection images of morula embryos (E2.5) cultured up to the early blastocyst stage (E3.5) with DMSO or inhibitors against DGAT1 and DGAT2 (DGATi; 25 µM and 50 µM) stained with BODIPY 493/503 (green), F-ACTIN (grey) and DAPI (blue). Scale bars, 50 µm.

(B) Number of BODIPY foci detected per embryo. Error bars, means ± s.e.m.; one-way ANOVA with post-hoc Dunnett's test (n=10-11) (see also Figure 5N).

(C) Scoring of the morphology of morula embryos cultured with DMSO or DGATi (25 µM and 50 µM) up to the early blastocyst stage.

(D) Schematic of image analysis approach using CellProfiler. DAPI-positive nuclei were segmented in Gaussian-smoothed single z-sections. ZSCAN4 signal intensity was quantified within segmented nuclei (see also Figure 5O).

(E) Representative images of blastocysts with DMSO or DGATi (25 µM) stained with ZSCAN4 (pink), BODIPY 493/503 (green), F-ACTIN (grey) and DAPI (blue). Scale bars, 50 µm.

### FIGURE S6

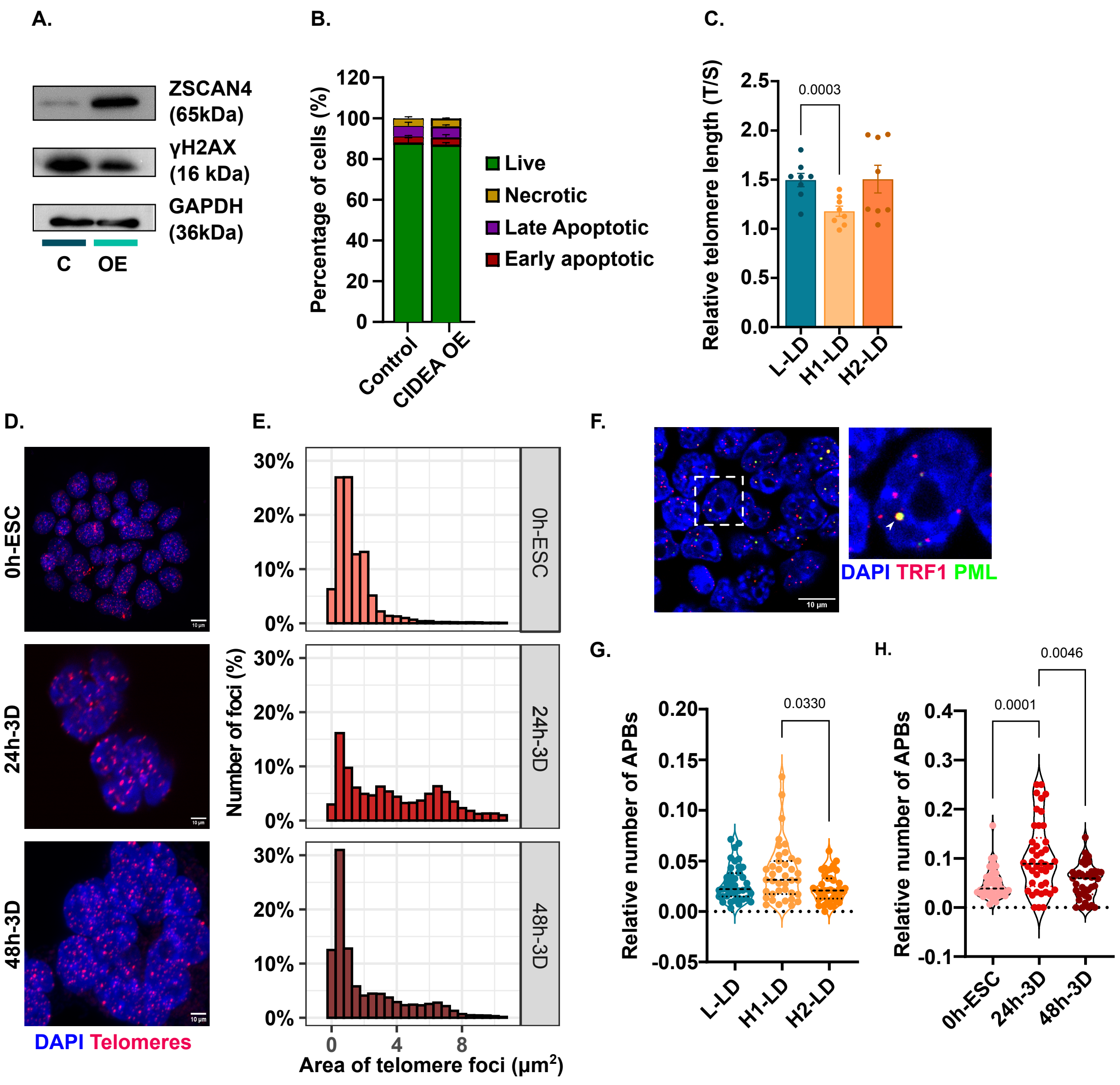
A.
ZSCAN4 (65kDa)
γH2AX (16 kDa)
GAPDH (36kDa)
C
OE
B.
Percentage of cells (%)
Live
Necrotic
Late Apoptotic
Early apoptotic
Control
CIDEA OE
C.
Relative telomere length (T/S)
0.0003
L-LD
H1-LD
H2-LD
D.
0h-ESC
24h-3D
48h-3D
DAPI Telomeres
E.
Number of foci (%)
Area of telomere foci (μm2)
F.
DAPI TRF1 PML
10 μm
G.
Relative number of APBs
0.0330
H.
0.0001
0.0046
0h-ESC
24h-3D
48h-3D

**Figure S6: Telomere homeostasis in ESC-based models prior to and upon differentiation, related to Figure 6.**
(A) Representative western blots of ZSCAN4 and γH2AX (pSer139) with GAPDH loading control in CIDEA OE (OE) and control (C) ESCs.
(B) Measurement of apoptosis indices in CIDEA OE and control ESCs using flow cytometry-based measurements of Annexin-V and propidium iodine (n=3).
(C) Relative telomere lengths of representative L-LD (8), H1-LD (4) and H2-LD (11) clones as measured by telomere qPCR, shown as a ratio of telomere to single-copy gene (T/S). Error bars, means ± s.e.m.; paired two-tailed t-test with Welch's correction (n=8). (D) Representative images of Telo-FISH, under non-denaturing conditions, using the TelC-Cy3 probe (red) and DAPI (blue) in undifferentiated ESCs (0 hour) and ESC-derived 3D spheroids at different timepoints of differentiation (24 and 48 hours).
(E) Frequency distributions of the area ($\mu m^2$) of telomeric foci at each timepoint (n=90 colonies/spheroids across 3 independent experiments).
(F) Representative image of ALT-associated PML bodies (APBs) as visualized by co-localization of PML (green) to telomeres (TRF1; red) within DAPI-stained nuclei (blue). Scale bars, 10 µm.
(G,H) Quantification of number of APBs per nucleus, normalised to total PML in (G) L-LD (8), H1-LD (4) and H2-LD (11) clones, and in (H) ESC-derived 3D spheroids at different timepoints of differentiation. ANOVA with Dunnett's post-hoc test (n>40 colonies/spheroids across 3-4 independent experiments).
